# Human Deep Cortical Neurons Promote Regeneration and Recovery After Cervical Spinal Cord Injury

**DOI:** 10.1101/2021.08.11.455948

**Authors:** Vanessa M. Doulames, James Weimann, Giles W. Plant

**Affiliations:** Department of Neurosurgery, Stanford University School of Medicine, Stanford, CA 94305, USA

## Abstract

Cervical spinal cord injuries (SCI) sever and permanently disrupt sensorimotor neural circuitry. Restoring connectivity within the damaged circuitry is critical to improving function. Herein we report robust regeneration of severed neural circuitry in a rat SCI model following transplantation of human induced pluripotent cells differentiated towards a deep cortical neuron lineage (iPSC-DCNs). *In vivo*, iPSC-DCNs: (1) integrated within the damaged cord and extended axons to caudal targets, (2) reversed SCI pathophysiology, (3) promoted robust regeneration of severed host supraspinal neural tracts, (4) and improved sensorimotor function. The results herein represent a significant paradigm shift in anatomical and functional outcomes over current preclinical/clinical models and demonstrates the survival and efficacy of human stem cell-derived cortical neurons in a SCI.

## Introduction

Cervical spinal cord injuries (SCI) permanently disrupt the neural tracts responsible for sensorimotor function^1–7^. Regeneration of these SCI-severed supraspinal tracts across injured tissue remains the largest obstacle impeding recovery. Therefore, we sought to intraspinally transplant a cellular population that could support and augment host axonal regeneration through the injury and allow for de novo sensorimotor circuit formation (**Fig. 1A**).

**Figure 1.**
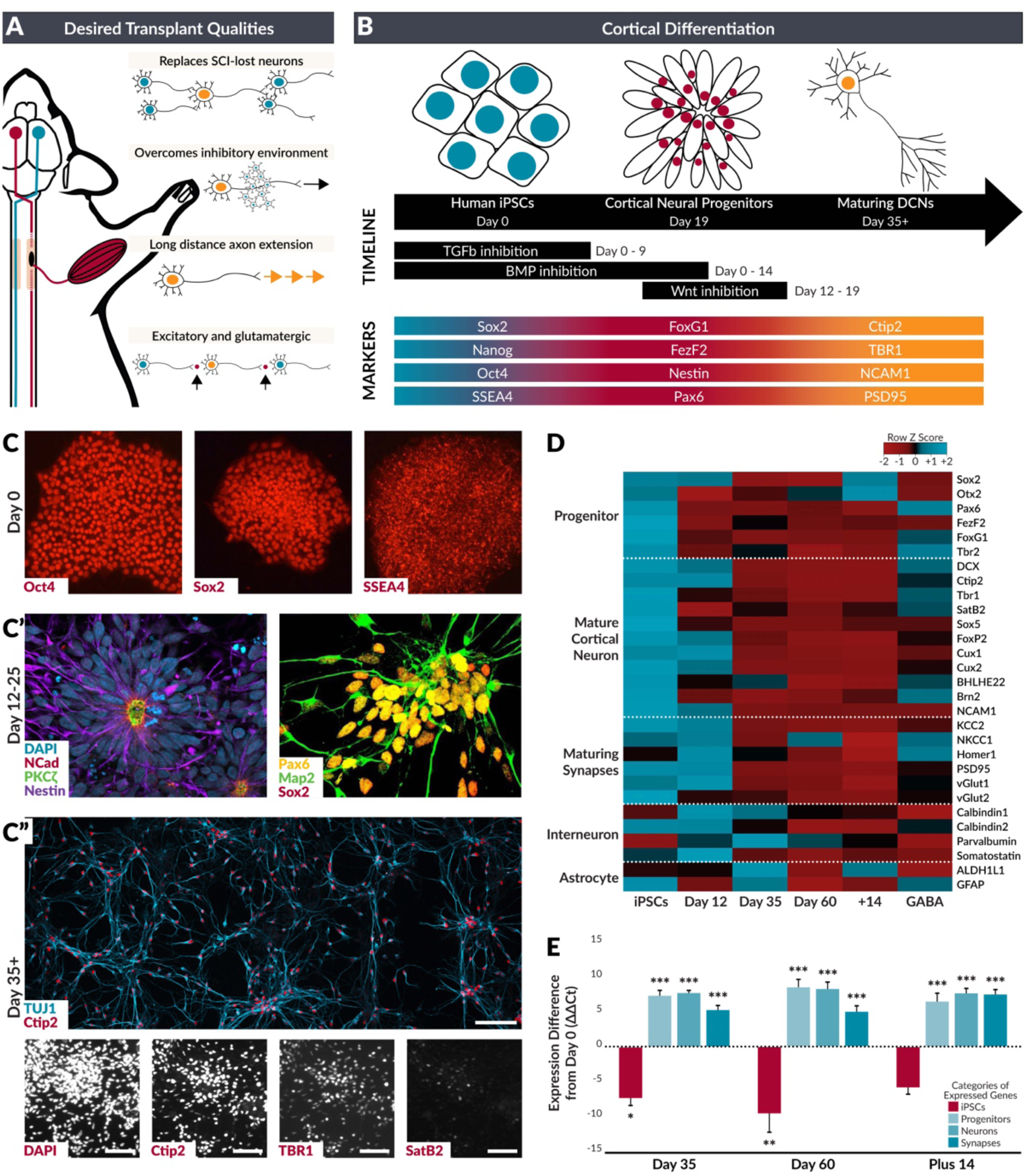
*In vitro* characterization of iPSC-DCNs. **A.** Schematic depicting characteristics that may be important when designing cellular therapies for SCI. **B.** Graphical description of the directed differentiation protocol used to derive a deep cortical neuron phenotype from induced pluripotent stem cells. **C.** Confirmation of the phenotypic identity of human iPSCs by immunohistochemical expression of proteins characteristic of stem cells (Day 0; Oct4, Sox2, SSEA4), neural 5progenitors (**C’**. Day 129-25; NCad, Nestin,PKCzeta, Pax6, Map2, Sox2), and maturing layer V/VI deep cortical neurons (**C”.** Day 35+; Ctip2, TBR1, and reduced expression of SatB2). Scalebars = 50um. **D.** Heatmap generated from qRTPCR analysis of iPSC cultures at different ages across a developmental timeline. Z scores calculated from ΔCt values. **E.** ΔΔCt values of groups of genes (iPSC: Oct4, Nanog; Progenitors: Sox2, Otx2, PAX6, FezF2, FoxG1, Tbr2; Neurons: NCAM1, CTIP2, Tbr1, SatB2, Sox5, FoxP2, Cux1, Cux2, BHLHE22, Brn2; Synapses: KCC2, NKCC1, Homer1, PSD95, vGlut1, vGlut2) in iPSC-derived cultures at different developmental ages were compared to the average ΔΔCt values of the Day 0 cohort. Averages are represented as columns ± SEM. Holm Method; *P ≤ 0.05, **P ≤ 0.01, ***P ≤ 0.001.

Normal sensorimotor function is mediated by cortical neurons with cell bodies that reside in the deep layers of the sensorimotor cortex. These developmentally-defined cell types extend axons in long tracts to directly innervate neurons that control function in distal locations of the spinal cord. In development, these long projections also provide permissive substrates for the extension of later developing axons from the cortex to distal targets in the spinal cord, and we hypothesized that these cell types might provide a bridging substrate for re-growth in the context of SCI.

Herein, human induced pluripotent stem cells were differentiated into a highly enriched deep cortical neuron lineage (iPSC-DCNs) and directly transplanted into a rat model of acute cervical SCI (**Fig. S1**). Quantitative analyses demonstrate that human iPSC-DCNs safely and stably integrate within the injured cervical spinal cord and maintain their phenotypic identity for at least one year post transplantation. The iPSC-DCN transplant attenuates cervical SCI pathology, elicits robust host axonal regeneration that bridges the lesion to innervate distal spinal segments, and provides definitive measurable improvements in sensorimotor function.

## Results

### Human iPSCs can be driven towards a deep cortical neuron phenotype

*In vitro*, both embryonic and induced pluripotent stem cells can undergo directed differentiation that recapitulates the temporal expression of fate-specifying transcription factors during human cortical neurogenesis^8, 9^. To characterize cellular identity prior to transplantation, three human iPSC lines were differentiated towards a cortical neuron fate using an adaptation of established protocols (**Fig. 1B**). Prior to differentiation, human iPSCs demonstrate appropriate morphology and expression of pluripotency markers (**Fig. 1C-E**). Within 10 days of neural induction, cells radially organize into rosette structures, a characteristic fundamentally inherent to neuroepithelium (**Fig. 1C’**)^10^. By Day 35 and beyond, Tbr1 (post-mitotic corticothalamic projecting neurons of Layer VI) and Ctip2 (subcerebral projecting neurons of Layer V) are expressed (**Fig. 1C”**) while SatB2 expression is reduced, indicating a progression from corticogenesis to a mature, subcortical projecting neuron identity^11^. This was accompanied by the expression of excitatory glutamatergic pre- and postsynaptic markers PSD95, Homer1, and vGlut1/2 (**Fig. 1D-E**). qRT-PCR confirmed the expression of additional genes and a strong enrichment for neurons with a terminal deep layer cortical neuron identity as cultures differentiate (**Fig. 1D-E**). Collectively, this suggests that human iPSCs are able to terminally differentiate into spontaneously active deep layer cortical projection neurons following previously established *in vitro* and *vivo* time courses and expression patterns^8–12^.

### iPSC-DCNs survive transplantation and extend axons beyond the SCI border

There is little evidence supporting the survival and integration of maturing neurons within the adult injured central nervous system. Therefore, we first examined iPSC-DCN survival, distribution, axonal outgrowth, and stability at 12 weeks and 1 year post transplantation. Of the rats transplanted with human iPSC-DCNs, 90% had evidence of transplant survival (**Fig. 2**). Transplanting iPSC-DCNs with supportive growth factors^13^, did not significantly affect transplant survival or volume (**Fig. 2B**). Approximately half of quantified SC101^+^ human iPSC-DCNs were found at the transplant epicenter (**Fig. 2C-D**) while 97% of the transplanted cells were located within 3 mm of the epicenter. Spinal cord volumes occupied by iPSC-DCNs and the rostral-caudal distribution of cells were similar across animals (**Fig. S2**). iPSC-DCN cell bodies were not detected in the contralateral side of the spinal cord, or any location within the brain stem and brain. Human SC121^+^ iPSC-DCN axons extended well beyond the lesion scar for over 1.5cm from the transplant epicenter to reach the thoracic spinal cord.

**Figure 2.**
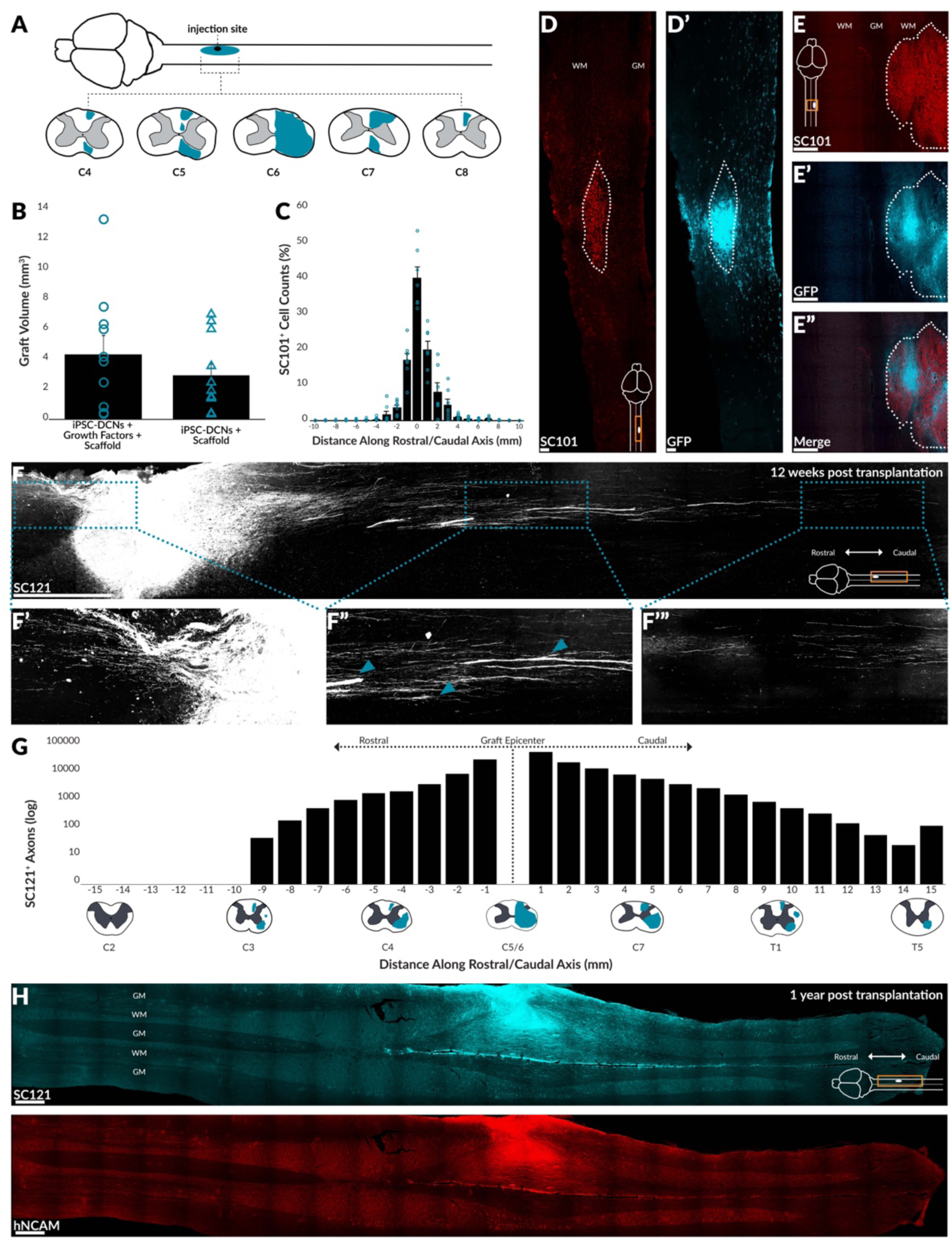
Human iPSC-DCNs survive transplantation, show self-limiting migration into the lesion penumbra, and extend axons long distances in vivo. **A.** Schematic depicting the spread of SC101^+^ grafted human cells spinal segments after 12 weeks. **B.** Graft volume was determined by quantifying the area in which SC101^+^ cells were located. Averages (n=10 per cohort) are represented as columns ± SEM with individual values overlayed (blue, circles and triangles). **C.** SC101^+^ human iPSC-DCNs were quantified in 1 mm increments from the graft epicenter after 12 weeks. Averages are represented as columns ± SEM with individual values overlayed (blue circles). **D-E.** SC101^+^/GFP^+^ grafted iPSC-DCNs (dotted outline) in 2 representative rats at 12 weeks post transplantation. WM = white matter, GM = gray matter. Scale bars = 100 µm and 500 µm, respectively. **F.** Qualitative image depicting successful iPSC-DCN engraftment at 12 weeks post transplantation. Arrows highlight SC121^+^ fiber bundles (**F”**). Scale bar = 1 mm. **G.** SC121^+^ labeled axons of transplanted iPSC-DCNs in a subset of rats (n=8). Schematic in X-axis corresponds with representative locations of quantified SC121^+^ human axons. **H.** SC121^+^/NCAM^+^ iPSC-DCN engraftment at 52 weeks post transplantation. WM = white matter, GM = gray matter. Scale bars = 1 mm.

Axons also extended 9 mm rostral from the transplant epicenter, bridging the SCI lesion to reach the C3 segment of the cervical spinal cord (**Fig. 2F-G**). Axons predominantly localized within the right dorsal and lateral funiculi (corresponding with corticospinal tract projections in the rat) and within the ventral motor neuron pools of caudal segments (**Fig. 2G**). While fiber densities varied, the overall length of rostral and caudal projections was consistent between animals (**Fig. S3**). Transplanted rats euthanized 1 year post transplantation confirmed continued survival with similar soma and axonal distribution as seen at 12 weeks (**Fig 2H**). Collectively this suggests that iPSC-DCNs (1) are capable of robust survival without additional trophic support, (2) exhibit self-limiting migration/dispersion^14, 15^, (3) and are able to integrate into the adult injured spinal cord despite the purported axon-inhibitory milieu of the SCI site.

### iPSC-DCNs stably maintain a deep cortical neuron phenotype in vivo

The SCI microenvironment influences the final phenotypic identity of transplanted cells not yet committed to a specific fate. Therefore, we have addressed the important question of whether the SCI microenvironment would influence the final phenotypic identity of transplanted, fate-restricted iPSC- DCNs (**Fig. 3A**).

**Figure 3.**
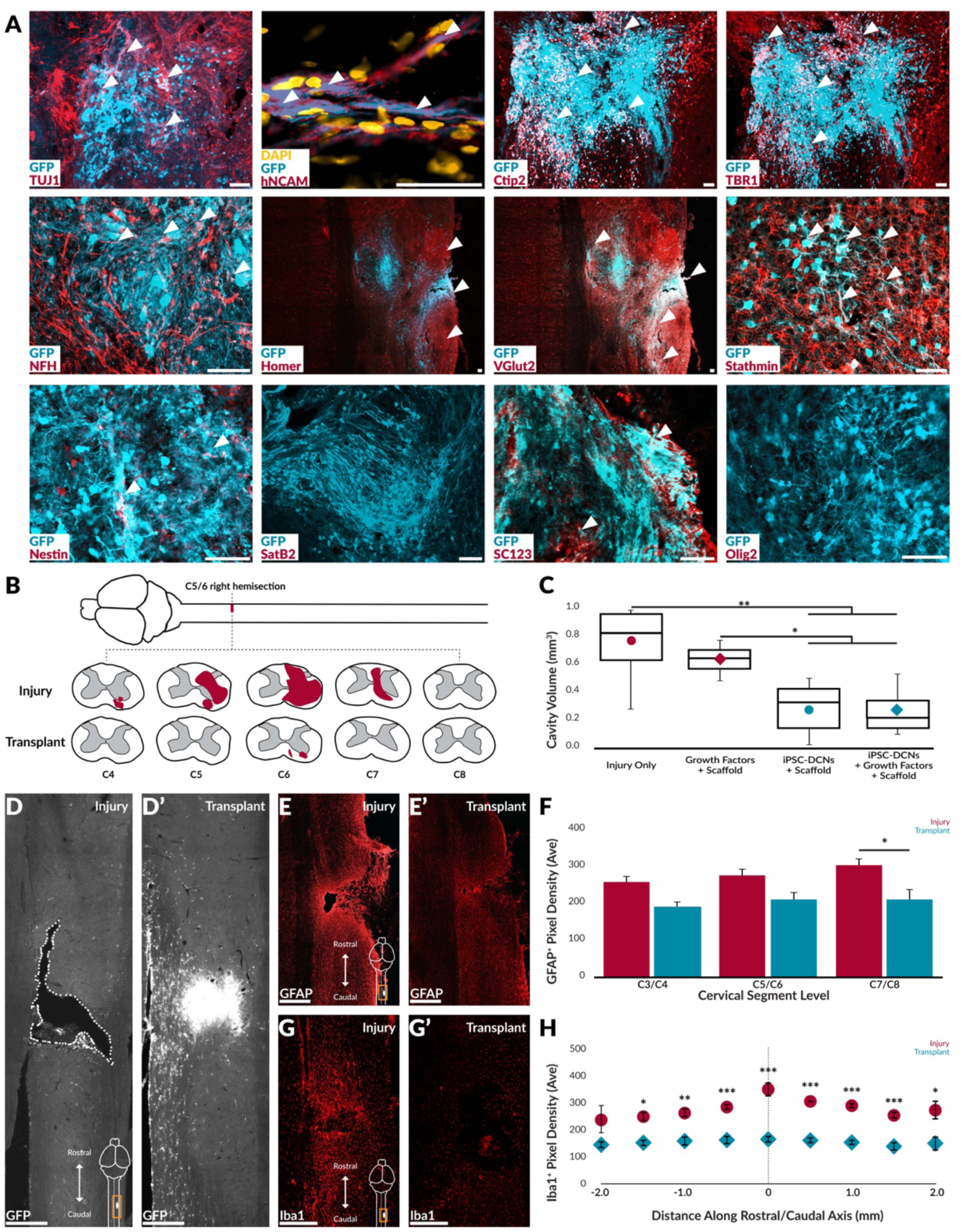
Human iPSC-DCNs maintain their deep cortical identity *in vivo* and reduce cavitation and inflammation. **A.** Grafted GFP^+^ iPSC-DCNs are TUJ1^+^, human NCAM^+^, NFH^+^; Ctip2^+^, TBR1^+^, Homer^+^ and VGlut 2^+^. Low numbers of GFP^+^ iPSC-DCNs are Stathmin^+^, Nestin^+^, SC123^+^, but not Satb2^+^ or Olig2^+^. Scale bars = 50 µm. **B.** Shaded areas show cavitation from C4-C8 in cords of transplanted and control rats. **C.** Lesion cavity volume results are represented as box and whisker plots to visualize distribution; (red = no cells, blue = cells). Tukey HSD Test; \**P ≤ 0.05*, \*\**P ≤ 0.01*. N=11, 3, 7, and 16 respectively. **D.** GFP^+^ human iPSC-DCNs (**D’**) fill the lesion cavity (dotted outline) (**D**). Scale bars = 500 µm. **E-F.** iPSC-DCNs (n=8) reduce GFAP^+^ astrogliosis compared to injury controls (n=6) at rostral, lesion, and caudal segments. Fluorescence intensity averages are represented as columns ± SEM. Tukey HSD Test; \**P ≤ 0.05*. Scale bars = 500 µm. **G-H.** After 12 weeks, iPSC-DCN transplantation (n=8) reduces Iba1^+^ microglia/macrophage presence compared to injury controls (n=6). Fluorescence intensity averages are represented as points ± SEM. Welch’s T-Test; \**P ≤ 0.05*, \*\**P ≤ 0.01*, \*\*\**P ≤ 0.001*. Scale bars = 500 µm.

At 12 weeks post transplantation, GFP^+^ iPSC-DCNs were positive for mature neuronal markers (TUJ1, human NCAM, and NFH), deep layer V/VI cortical-specific markers (Ctip2, TBR1), and excitatory, glutamatergic synaptic markers (Homer, VGlut2) (**Fig. S4**). At this timepoint, the transplant is also positive for neural progenitor markers Nestin and Stathmin (specific to deep Layer V/VI cortical maturing neurons)^16^ but not SatB2. A small component of transplanted cells were SC123^+^ (human astrocytic marker). There was no immunohistochemical evidence that transplanted iPSC-DCNs returned to a pluripotent state (Pax6 and Sox2), differentiated towards an oligodendroglial phenotype (Olig1), or exhibited unregulated growth or tumor formation. Consistent with the literature, this suggests that transplanting a fate-restricted population of cells overcomes the SCI microenvironment’s influence on differentiated identity.

### Transplantation ameliorates classic SCI pathology

SCI results in a cystic cavity within the margins of a glial scar, formed by reactive astrocytes and their secretion products. Regeneration through the cavity is diminished by the lack of available tissue scaffolding for axonal growth. Resident Iba1^+^ microglia and monocyte-derived macrophages further shape host immune and inflammatory responses over the lifetime of the animal. This is via the secretion of molecules, phagocytosis, and their direct influence on astrocytes, oligodendrocytes, and demyelination. Therefore, we asked whether iPSC-DCN transplantation could modulate these hallmarks of SCI pathology. This model resulted in a chronic lesion cavity that could extend as far as four segments post injury (**Fig. 3B-D**), which was significantly reduced by iPSC-DCN transplantation with and without exogenous growth factors. In the absence of cells, injecting fibrinogen/thrombin matrix with and without growth factors did not reduce cavitation.

iPSC-DCN transplantation significantly reduced GFAP^+^ reactive astrogliosis caudal to the lesion and modestly rostral and within the lesion, compared to injured controls (**Fig. 3E-F**). Despite reduction, astrocytes are still present in the transplant zone, along with inhibitory factors such as chondroitin sulfate proteoglycans (CS56) (**Fig. S5**). iPSC-DCN transplantation also significantly reduced Iba1^+^ microglia/macrophage presence compared to injured controls (**Fig. 3G-H**). In injured control rats, Iba1^+^ presence is strongest at the lesion epicenter and diminishes for 2 mm in either direction. Conversely, transplanted rats show no variation over distance.

Collectively, this suggests that (1) transplanted cells are required to fill the lesion cavity and that exogenous delivery of growth factors in an extracellular matrix is not sufficient to impact cavitation, and (2) iPSC-DCNs are capable of ignoring or overcoming astrogliosis/inflammation-mediated inhibitory cues, which may contribute to the ability of axons to extend through and beyond the scarred border into intact spinal tissue.

### Transplantation leads to corticospinal tract regeneration and reinnervation

The corticospinal tract is responsible for voluntary sensorimotor function and does not regenerate post injury in higher adult vertebrates; this has been reported to be a response to inhibitory cues within the environment rather than an innate inability of the severed axons to regrow^17, 18^.Recent evidence has suggested descending corticospinal tract and serotonergic (5HT) tract axons can regenerate through chronic scars of the CNS if given appropriate positive growth assistance.^19^ We therefore asked whether iPSC-DCN transplantation could overcome the inhibitory SCI milieu and induce regeneration of severed corticospinal tract axons. This was examined by anterogradely tracing the projections of host corticospinal neurons using adeno-associated viruses (AAVs) expressing mCherry (contralateral motor cortex) or GFP (ipsilateral motor cortex). In injured control rats, contralateral projecting mCherry^+^ axons did not extend beyond the rostral lesion border. In contrast, iPSC-DCN transplanted rats demonstrated robust regeneration of severed contralateral projecting mCherry^+^ corticospinal axons (400% increase) traversing through the lesion into caudal segments of the spinal cord (**Fig. 4A**). These mCherry^+^ axons produced structures consistent with synaptic boutons within the target domains, suggesting successful synaptic innervation of caudal motor neuron pools (**Fig. 4E-F**). No difference was observed in the ipsilateral projecting GFP^+^ axons. In an additional cohort, transplanted rats retrogradely traced with Fluorogold (injected two segments caudal to the injury), had Fluorogold^+^ cell bodies within the contralateral motor and somatosensory cortices; this was not the case in injury control rats. Transplanted rats had a 12-fold higher number of Fluorogold^+^ cells within the contralateral motor cortex and a 4-fold higher number within the contralateral somatosensory cortex when compared to injured controls. These results provide evidence that host corticospinal neurons regenerated across the injury/transplant site to reinnervate caudal segments of the spinal cord (**Fig. 4G-J**). Collectively this suggests iPSC-DCN transplantation produces a highly growth supportive SCI microenvironment, which in turn promotes robust regeneration of severed corticospinal tract axons into and beyond the injury site.

**Figure 4.**
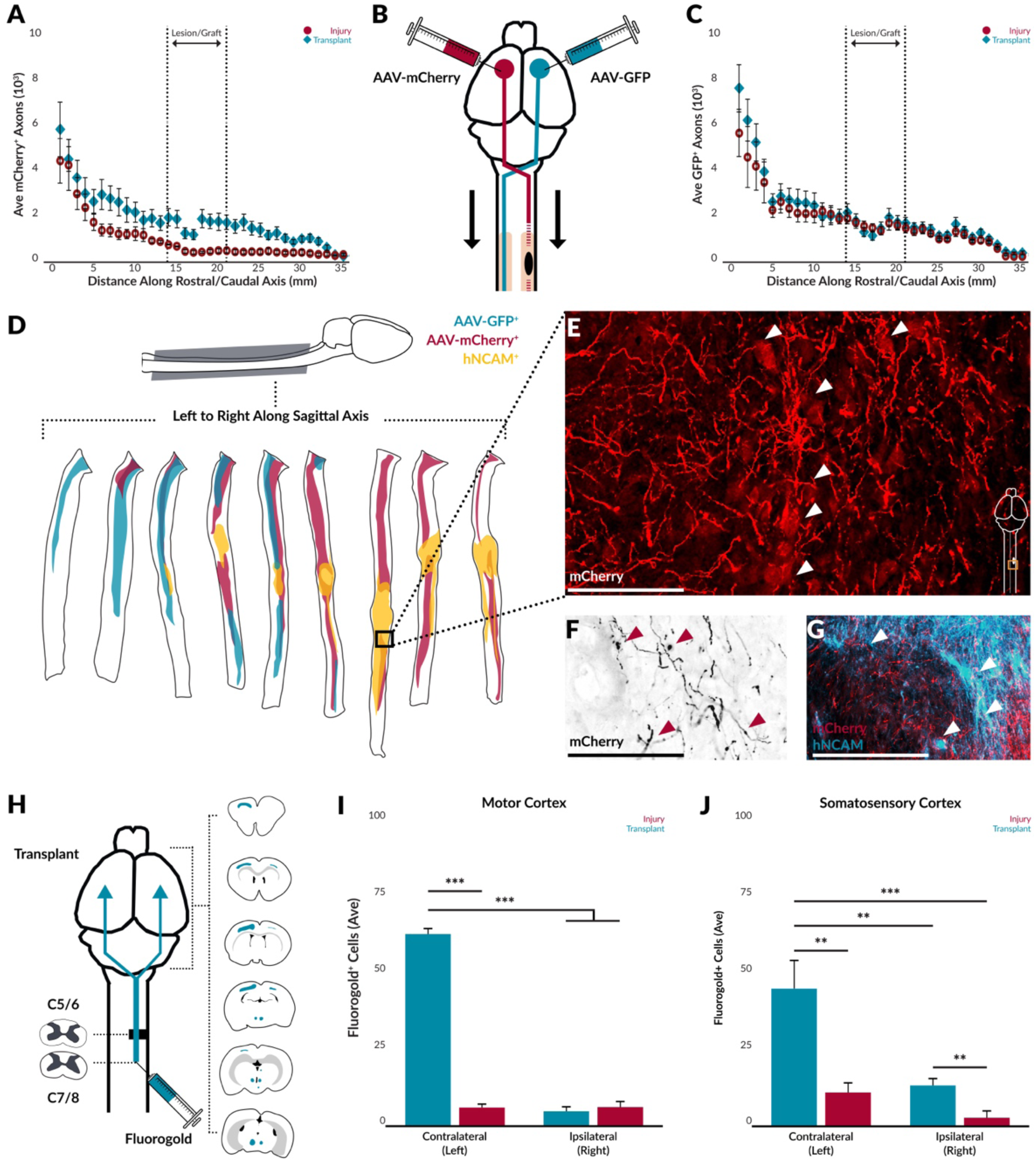
Human iPSC-DCN transplantation promotes corticospinal tract regeneration. **A-C.** GFP^+^ and mCherry^+^ were quantified in 1 mm increments from the brainstem down 3.5 cm in a subset of rats (n=10). Results are represented as the number of axons averaged across the cohort ± SEM. **D.** Spinal cord reconstruction of a representative transplanted rat in the sagittal plane to show the spatial distribution of GFP^+^ (blue) and mCherry^+^ (red) host corticospinal axons and hNCAM^+^ (yellow) transplanted iPSC-DCN axons. **E-G.** Insert images depicting host mCherry^+^ axons in motor neuron pools caudal to the lesion (**E**), with putative synaptic boutons (**F**), and in close proximity with transplanted hNCAM^+^ axons (**G**). Scale bars = 500 µm, 500 µm, and 250 µm respectively. **H.** Fluorogold^+^ cells were quantified within the brains of traced rats (n=10). Shaded blue areas indicate Fluorogold^+^ cells in the serial immunolabeled brain of a representative transplanted rat. **I-J.** Quantification of Fluorogold^+^ cells within the left and right motor and somatosensory cortices of injured and transplanted rats. Results are represented as the quantified number of cells averaged across the cohort ± SEM (red = injury control, blue = transplant). Tukey HSD Test; \*\**P ≤ 0.01*, \*\*\**P ≤ 0.001*.

### Transplantation improves regeneration and reinnervation of supraspinal tracts

Serotonergic (5HT) neurons have fiber tracts descending from the raphe nucleus that are distinct from glutamatergic fibers of host or transplanted iPSC-DCNs. In adult mammals post SCI, 5HT^+^ fibers sprout at the severed ends of descending axons but fail to bridge lesions in control animals or in lesions transplanted with other cell types^20, 21^. Furthermore, SCI-spared descending 5HT^+^ fibers and exogenous serotonin are reported to modulate functional recovery^22^. Therefore, we investigated if iPSC-DCN transplantation would influence regeneration of this tract and whether that would contribute to functional recovery. Consistent with previous reports, all rats displayed an increase in 5HT^+^ processes rostral and ipsilateral to the lesion^20^. In injured control rats, 5HT^+^ fluorescence was significantly reduced caudal to the lesion while in iPSC-DCN transplanted rats, 5HT^+^ fibers dramatically extend into the transplant and reinnervate the caudal spinal cord. We observed both host 5HT^+^ axons and transplanted GFP^+^ iPSC-DCN axons surrounding the same caudal motor neuron pools (**Fig. 5B**). Systemic blocking of serotonin using methysergide^23^ had no effect on behavioral function, suggesting 5HT^+^ reinnervation may impart a role in overall motor tone rather than task specific cervical-sensorimotor function (data not shown). Importantly, we found retrogradely labeled Fluorogold^+^ neurons in the raphe of transplanted rats (**Fig. 5E-F, Fig. S6**), supporting our data showing 5HT^++^ axons in immunostained spinal sections. Additionally, we quantified Fluorogold^+^ neurons in the brains and brainstems of injured and iPSC-DCN transplanted rats and found statistically significant numbers of FG labeled neurons in the lithoid, pararubral, red, vestibular, and reticular nuclei (**Fig. S6**). Collectively, these results demonstrate iPSC- DCN transplantation enhances the regenerative potential of endogenous supraspinal tracts severed by cervical SCI.

**Figure 5.**
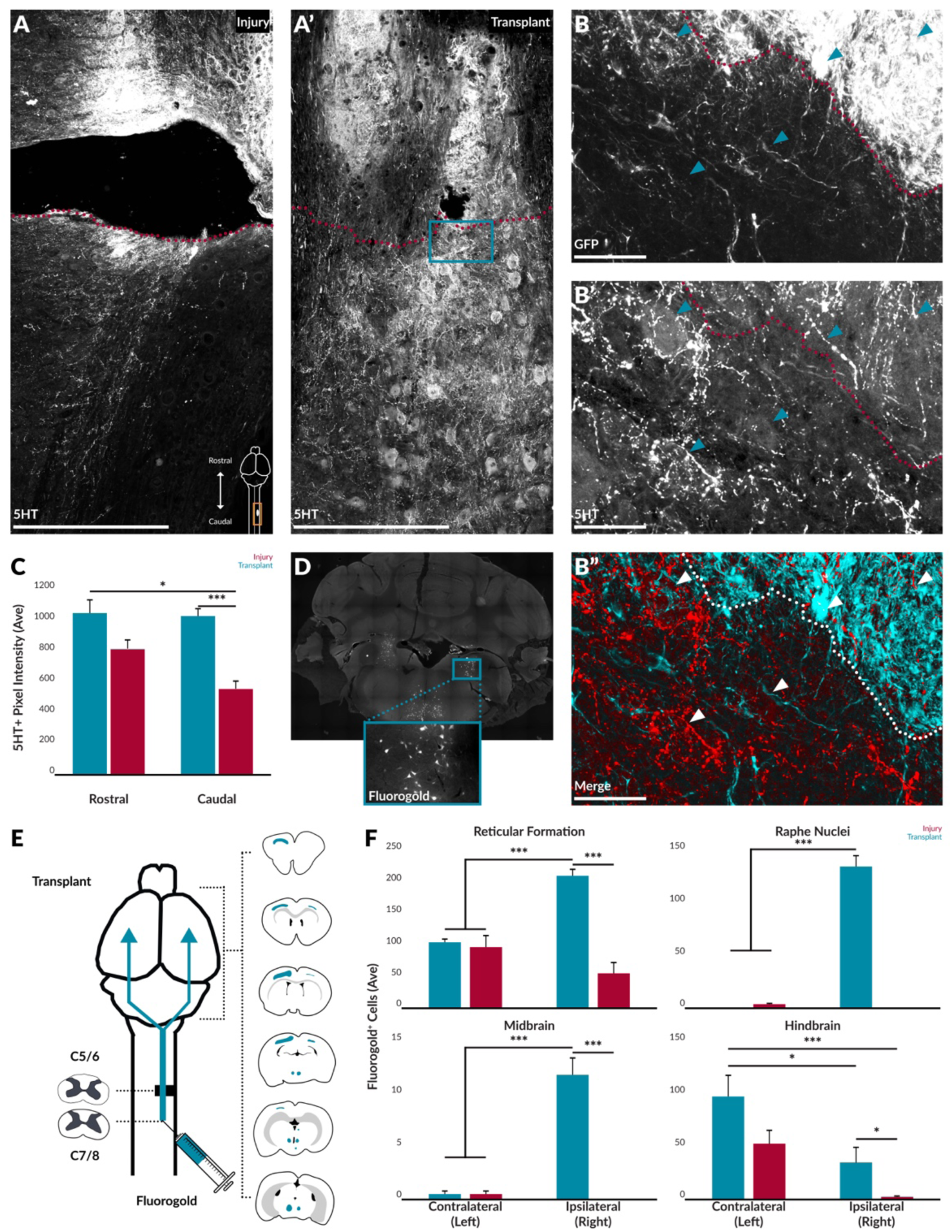
Human iPSC-DCN transplantation promotes regeneration of 5HT+ raphenergic axons. **A.** 5HT^+^ axons at the lesion epicenter in an injured and transplanted rat at 12 weeks post transplantation. Red dotted line delineates caudal border of the lesion. Scale bars = 500 µm. **B.** Magnified insert from **A’**. Dotted line delineates graft border (as evidenced by GFP^+^ labeling). Arrows illustrate motor neurons caudal to the lesion surrounded by transplanted GFP^+^ and host 5HT^+^ axons. Scale bars = 100 µm. **C.** 5HT^+^ fluorescence intensity was quantified within 2 segments rostral and caudal of the lesion epicenter. Fluorescence intensity averages are represented as columns ± SEM. Tukey HSD Test; \**P ≤ 0.05*, \*\*\**P ≤ 0.001*. **D.** Fluorogold^+^ labeling within the reticular formation of a representative transplanted rat at 12 weeks post transplantation. **E.** Shaded blue areas indicate the location and extent of Fluorogold^+^ cells in the serial immunolabeled brain of a representative transplanted rat. **F.** Quantification of Fluorogold^+^ cells within the brains of transplanted rats. Results are represented as the quantified number of axons averaged across the cohort ± SEM (red = injury control, blue = transplant). Tukey HSD Test; \**P ≤ 0.05*, \*\*\**P ≤ 0.001*.

### iPSC -DCN transplantation restores sensorimotor function across multiple behavioral assays

To determine if anatomical changes correlated with functional return, we used five different behavioral assays to track changes in SCI-lost sensorimotor function. To determine whether transplanted iPSC- DCNs were critical for improvements in function or supportive, we selectively depleted human iPSC- DCNs using diphtheria toxin (DPT) in a subset of rats.

Skilled walking was measured using the horizontal ladder rung test (**Fig. 6A**)^24^. Injury resulted in a 5- fold increase in total errors that was maintained over the study whereas transplanted rats recovered to pre-injury baselines. Post injury, the percentage of full slips rose 10-fold and partial slips rose 4-fold. All rats demonstrated a reduction in full slips over time yet never reached pre-injury levels. While injured control rats maintained this deficit in partial slips for the duration of the study, transplanted rats reached pre-injury levels by 10 weeks post transplantation. This suggests that motor control of forelimb lifting is only transiently affected by this injury (full slips). Forepaw sensation steadily declines, as there appears to be inappropriate sensory feedback leading to improper placement of the forepaw on the rung (partial slips). Transplanted rats return to baseline, further supporting our Fluorogold tract tracing data suggesting functional somatosensory innervation of the spinal cord caudal to the lesion.

**Figure 6.**
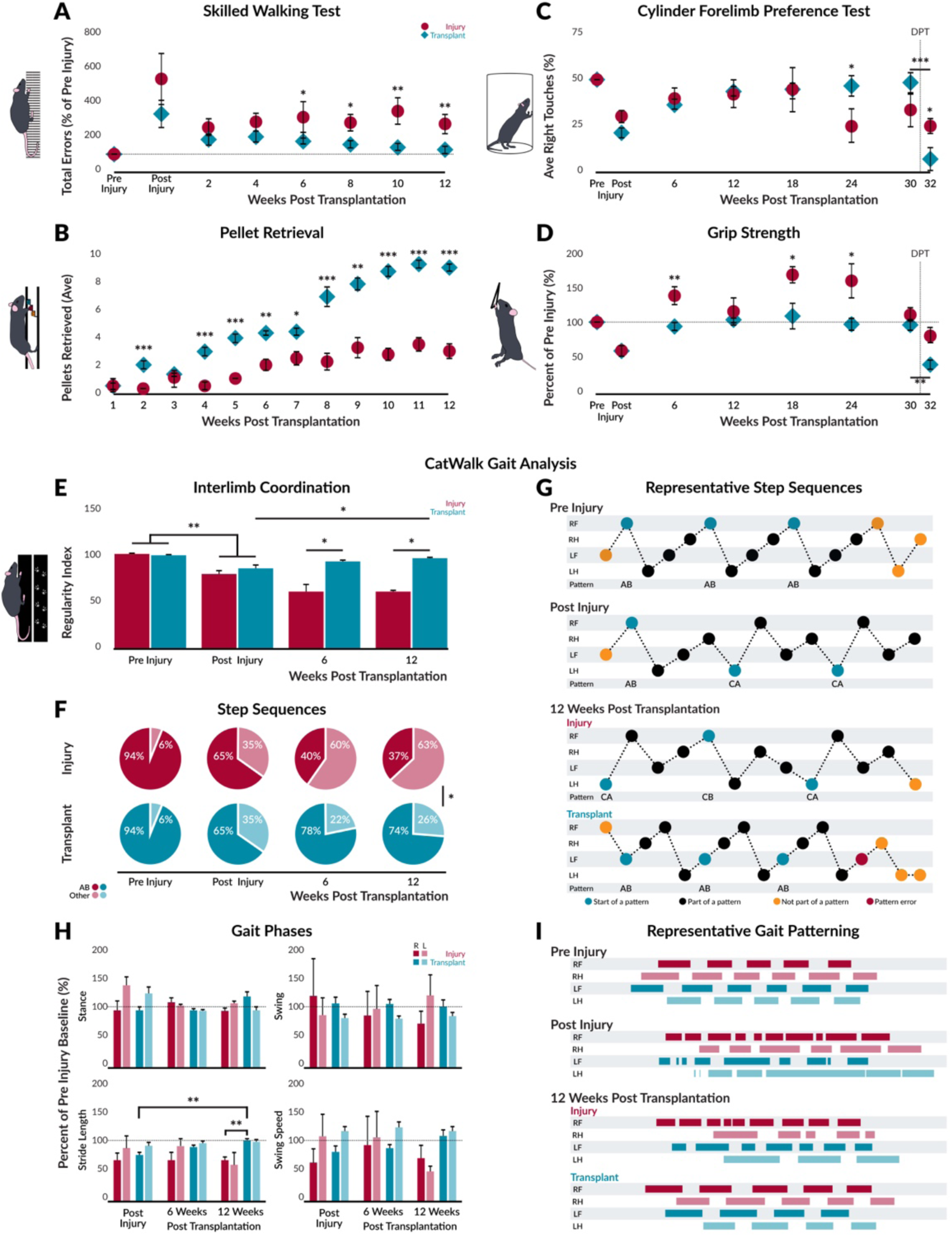
Human iPSC-DCN transplantation improves sensorimotor function. **A.** Skilled walking was measured using the ladder rung test (n=16). Total errors are represented as points ± SEM. **B.** Grasping was measured using the pellet retrieval test (n=15). Average pellets retrieved are represented as points ± SEM. **C.** The cylinder test measures symmetry in forepaw usage (n=68). The average percentage of right forepaw touches are represented as points ± SEM injury control. **D.** Grip strength (n=50) as a percentage of pre-injury is represented as points ± SEM. **E.** Interlimb coordination (n=16) was assessed by measuring regularity index (cohort mean ± SEM). **F.** Interlimb coordination (percentage of observed step sequence patterns (mean percent; dark shades = percentage of AB sequences, light shades = percentage of other sequences). **G.** Representative step sequences. **H.** Gait phases were assessed by measuring stance, stride length, swing, and swing speed of forelimbs (average percent of pre-injury ± SEM (dark shades = right forelimb, light shades = left forelimb)). **I.** Representative gait patterns. For all tests: red = injury control, blue = transplant; horizontal dotted line denotes pre-injury baseline; vertical dotted line denotes DPT administration. Welch’s T-Test; \**P ≤ 0.05*, \*\**P ≤ 0.01*, \*\*\**P ≤ 0.001*.

Functional grasping (the dynamic motion of extending the forelimb, grasping a food pellet, and bringing the pellet to the mouth) is severely impaired in the right forepaw after cervical injury and was measured using the pellet retrieval test (**Fig. 6B**). Post injury, rats are unable to grasp and retrieve food pellets. This deficit is maintained by injured control rats throughout the study while transplanted rats regain this ability and demonstrate a 3-fold increase in pellets retrieved.

The cylinder test measures symmetry in forepaw usage during vertical exploration (**Fig. 6C**). Pre-injury, usage of the forepaws is symmetrical but following injury, the percentage of right forepaw use drops 2- fold. In all rats, paw symmetry returns mid-study but this return to baseline is transient in injured control rats. In contrast, transplanted rats stably maintain equal paw usage for 30 weeks post transplantation, which is reversed 8-fold within 1 week of DPT ablation.

Grip strength is affected by injury as the motor neuron pools controlling the flexor and extensor muscles of the wrist are deafferented ^25–27^. This manifests as a substantial weakening (inability to grasp) or increase in grip strength (spastic inability to let go) (**Fig. 6D**). Injury initially reduced grip strength in all rats 2-fold. Injured control rats exhibited a transient increase in grip strength. In contrast, transplanted rats never exhibited this spastic strength and remained indistinguishable from pre-injury for 30 weeks post transplantation. However, within 1 week of DPT ablation, grip strength was reduced 2.5-fold from pre-injury.

Interlimb coordination during gait and locomotion was assessed by measuring regularity index (number of irregular paw placements relative to regular placements in a cruciate gait pattern) (**Fig. 6E**) and the percentage of step sequence patterns utilized by the rats (**Fig. 6F**). After injury, regularity index drops by 20% indicating dysregulation in gait. This is exacerbated in injured control rats throughout the study whereas transplanted rats return to baseline. Rodents have 6 observed step sequence patterns (**Fig. S8**) in which the AB sequence dominates. Injury results in a 1.5-fold reduction in usage of the AB sequence, which is exacerbated in injured control rats over the study. In injured rats, there was usually a combination of sequences used, further demonstrating dysregulation in coordination and stepping (**Fig. 6G**). In contrast, iPSC-DCN transplanted rats increased usage of the AB sequence resulting in more coordinated step patterning.

Stance, stride length, swing, and swing speed were analyzed to assess changes in gait phases (**Fig. 6H- I**). Injury resulted in a 25% decrease in right forelimb stride length. While this decrease was maintained by injured animals for the duration of the study, iPSC-DCN transplanted rats eventually return to baseline. No significant differences were observed in stance, swing, and swing speed across cohorts and experimental conditions.

## Discussion

Cervical SCI is devastating, yet survivable, resulting in lifelong disability for patients. In prior studies, transplanting mixed populations of neural progenitors has demonstrated variable efficacy in reducing and reversing SCI pathology. However, the SCI microenvironment influences transplanted progenitor survival, differentiation, and behavior ^28–30^; discerning how the mixed population will differentially respond to SCI or integrate and contribute to recovery becomes difficult to interpret. As such, surprisingly little is known as to how or why transplanted cells mediate regeneration and recovery ^31^. Experimentally transplanting a precisely defined, fate-restricted, and phenotypically stable population of cells provides the opportunity to learn if, and how, specific cellular identities differentially influence the injured spinal cord ^32–48^. Therefore, the objective of the present study was to test whether transplanting human iPSC-derived neurons patterned to acquire the identities of deep layer cortical projection neurons provide positive and lasting regenerative outcomes in a rat model of cervical SCI.

Cortical projection neurons residing in the deep layers of the sensorimotor cortex have a unique “fingerprint” of characteristics defining their phenotype and function; they exhibit gene expression patterns specific to their laminar fate, are capable of extensive axonal growth, and form glutamatergic excitatory synapses. Our results show that pre-patterning iPSCs into maturing DCNs allowed the transplanted cells to maintain this patterned phenotypic identity *in vivo*. Furthermore, transplanted iPSC- DCNs extended axons over long distances to terminate in caudal spinal motor neuron pools. The terminal axons had morphological structures resembling synaptic boutons, and were positive for glutamatergic synaptic markers. Finally, there was no evidence of hyperplasia or tumor formation for at least one year following transplant. Consistent with the literature ^32–48^, our results confirm that transplants driven towards a singular maturing phenotype reduces the strong instructive influence of the SCI microenvironment on differentiation and cell fate and instead allows transplants to retain a precisely defined and desired neuronal identity within the transplant.

The SCI microenvironment is known for its deleterious effects on host and transplanted cells; primary and secondary injury mechanisms contribute to large cysts surrounded by reactive cell types that create a physical barrier impeding severed axons from regenerating. iPSC-DCNs transplantation significantly reduced lesion cavity volume and reduced these cellular responses. Consistent with their patterned phenotype, transplanted iPSC-DCNs demonstrate robust axonal growth far beyond the lesion border, despite the presence of scarring that is repulsive to axonal regeneration. One possibility is that the human xenograft does not recognize rat-specific inhibitory cues within the injured spinal cord. This may contribute to the observed outcomes, but this does not adequately explain why host descending fibers are also able to transit the injury site in iPSC-DCN transplanted rats. A more intriguing possibility supported by the literature ^49^, is that iPSC-DCNs express transcriptional factors inherent to the phenotype that robustly overcome the inhibitory cues created following SCI and provide a growth-permissive bridging environment (similar to their developmental counterparts) that has functional benefits for the injured animals.

The corticospinal tract does not regenerate post injury in higher vertebrates and exhibits retrograde Wallerian degeneration; this is not an innate inability of the severed axons to grow but rather a response to inhibitory cues within the lesion environment ^17, 18^. Therefore, the robust regeneration of severed anterograde labeled host corticospinal tract axons we observed strengthens our hypothesis that iPSC- DCN transplants positively alter the SCI microenvironment, and allow bridging and regeneration of severed corticospinal axons to appropriate caudal targets.

In the literature, exogenous neurotrophin administration augments serotonergic regeneration and in high cervical injuries, 5HT or 5HT agonists promote neuroplasticity of injured phrenic circuits ^50, 51^. Additionally, serotonergic regeneration is well-established as predictive of functional recovery following SCI in lower spine segments ^52–54^. In transplanted rats, the iPSC-DCN transplants were robustly innervated with host 5HT^+^ processes, which exited the lesion to surround caudal deafferented motor neuron pools in tandem with human iPSC-DCN axons. Collectively, this further supports the hypothesis that iPSC-DCNs are modulating the permissiveness of the lesion in a manner that encourages regeneration of supraspinal SCI-severed axons and contributes to sensorimotor recovery.

In rodents, mid-cervical SCI deafferents the spinal motor neuron pools responsible for forelimb function. While lesions in lower spinal segments are associated with transient or minor sensorimotor deficits, cervical injury permanently impairs fine motor function ^24^. We observed significant sensorimotor recovery in seven parameters over five different behavioral assays. iPSC-DCN transplantation was associated with significant improvements in skilled walking, dynamic grasping, gross motor function of the upper forelimbs, grip strength and dexterity, and regulated over ground gait and locomotion. Furthermore, selective ablation of the human iPSC-DCN transplant using diphtheria toxin reversed any functional gains while injured control animals experienced no significant effect. This suggests that, rather than functioning simply as a substrate in which host axons could regenerate through, transplanted iPSC-DCNs may form novel circuits with host tracts culminating in a return of sensorimotor function.

Looking ahead, the data presented here establishes an important foundation to begin studying novel mechanisms of sensorimotor circuit formation post injury and transplantation, which will allow for greater control and fine-tuning within this paradigm. These results show that rostralized deep layer projection neurons can be used to successfully attenuate the functional deficits following cervical SCI, and establishes this phenotype as a strong therapeutic contender for cellular therapy.

## Methods

### Study Design

#### Sample Size

For the first replicate, sample size was based on historical cohort sizes found in similar studies in the literature. Based on the results from the initial replicate, a power analysis was performed using Sigma Plot v14.5 (Systat Software, Inc., San Jose, CA) for each subsequent replicate to calculate an appropriate sample size for further testing.

#### Replicates

Four replicates were performed in which behavioral and immunohistochemical data were collected. Axonal tracing experiments were performed twice, and transplant ablation was performed once. The number and composition of replicates were based on the experiments necessary to assess the collective effects of this transplantation strategy. The therapeutic results of this strategy have been substantiated and remain consistent under a range of conditions, including: gender and identity of transplanted cell source, different surgeons, and spinal cord injury models with varying locations and severities.

#### Data Collection

Defined rules for stopping data collection was outlined in advance based on set humane and study endpoints. Data exclusion criteria was determined in advance; data from individual rats were excluded if: i) there was early mortality due to complications, ii) the hemisection injury was inappropriate (too much or too little severed based on *post-mortem* tissue reconstructions) or iii) if there was no evidence of iPSC-DCN transplantation (assessed via GFP or human markers SC121 and SC101 immunofluorescence). The collective number of excluded rats was reported. All outliers were reported and included in the data analysis unless the rat fell under the exclusion criteria listed above.

#### Study Endpoints

Study endpoints were selected in advance; 12 weeks post transplantation allows for transplanted cells to stabilize and integrate within the host tissue, while 32 and 52 weeks post transplantation are meant to qualitatively represent long-term stability of the transplant and effects.

### iPSC-DCNs

#### Human iPSC Care and Culture

iPSC lines 1815c7, 32FSCNOCI, and HUF5 were used. iPSCs were plated in 6 well plates (Corning Inc., Corning, NY) that had been coated with Matrigel Basement Membrane Matrix (BD, Franklin Lakes, NJ) at 4°C overnight. They were maintained on an E8 Media cocktail changed daily consisting of: DMEM/F12 with Glutamine and HEPES (Invitrogen, Thermo Fisher Scientific, Waltham, MA), l-ascorbic acid 2-phosphate (final concentration 64 ug/mL; Sigma-Aldrich, St. Louis, MO), sodium selenite (final concentration 14 ng/mL; Sigma-Aldrich), transferring (final concentration 10.7 ug/mL; Sigma-Aldrich), insulin (final concentration 20 ug/mL; Invitrogen), FGF2 (final concentration 100 ng/mL; PeproTech, Inc., Rocky Hill, NJ), and TGFb1 (final concentration 2 ng/mL; PeproTech). Morphology was monitored daily for cobblestone appearance, uniform and tightly packed cells, and clearly defined borders. Cultures were passaged prior to changes in morphology consistent with large colonies. Remnant media was aspirated, and the culture was gently rinsed with PBS without calcium or magnesium. EDTA (0.5 mM in PBS) was added to the culture as a chelating agent for 5 minutes at room temperature. Once changes in morphology started to occur (edges of culture curling up), EDTA was aspirated and E8 media was used to remove adherent cells from the plate where it was then gently triturated prior to replating. Rho-associated protein kinase inhibitor (3 uM; 1293823; PeproTech) was added to the media for 1 day post replating.

#### Directed Deep Cortical Differentiation of iPSC Lines

iPSCs were maintained in E8 media until the culture was about 80% confluent with standard, uniform morphology. At this point, E8 media was aspirated and replaced with Differentiation Media (all components: Gibco, Thermo Fisher Scientific, Waltham, MA) consisting of: DMEM/F12 (48.25%), Neurobasal-A (48.25%), B-27 supplement sans serum (1%), N-2 supplement (0.5%), GlutaMAX supplement (1%), and MEM non-essential amino acids solution (1%), designed to stimulate cortical neurogenesis via the removal of FGF2.^55^ TGFb and BMP signaling were blocked using small molecules LDN-193189 (500 nM, Day 0-19; Stemgent, Cambridge, MA) and SB431542 (10 uM, Day 0-9; Stemgent) to drive the cultures towards a neuroepithelium fate. LDN-193189 is a cell-permeable BMP inhibitor of ALK2 and ALK3 while SB431542 inhibits ALK4, ALK5, and ALK7. On Day 12, the differentiating cultures (now dorsal neural progenitors) were dissociated onto poly-d-lysine (50 ug/mL diluted in 0.1 M Borate Buffer, pH 8.6) and laminin (10 ug/mL; Roche, Basel, Switzerland) coated 6 well plastic plate. The culture is then rostrally shifted towards a dorsal telencephalic fate via Wnt inhibition using small molecules Wnt-C59 (5 nM, Day 12-19; Stemgent). Wnt-C59 prevents palmitylation of Wnt proteins by porcupine (Porcn) and blocks Wnt secretion and activity. After 35 days of differentiation, 80% of the cells are maturing deep layer cortical neurons and the population is papain-dissociated using a commercial protocol and system (Worthington Biochemical Corporation, Lakewood, NJ) for transplantation ^56^.

#### Quantitative RT-PCR

For quantitative RT-PCR, total RNA was isolated from iPSC cultures driven towards glutamatergic deep cortical neurons or GABAergic neurons at time points along the differentiation protocol spectrum (Days 0, 12, 35, 60, and 35+14; Trizol, Sigma-Aldrich). Total RNA was reverse transcribed and used for quantitative RT-PCR with primers specific to *Sox2, Otx2, Pax6, FezF2, FoxG1, Tbr2, DCX, Ctip2, Tbr1, SatB2, Sox5, FoxP2, Cux1, Cux2, BHLHE22, Brn2, NCAM1, KCC2, NKCC1, Homer1, PSD95, vGlut1, vGlut2, Calbindin1, Calbindin2, Parvalbumin, Somatostatin, ALDH1L1*, and *GFAP* using the Applied Biosystems™ ABI PRISM™ 7000 Sequence Detection System (Applied Biosystems, ThermoFisher Scientific, Waltham, MA) and Rotor-Gene Q (Qiagen, Hilden, Germany) systems. Negative controls were used to test for contamination and melting curves were used to evaluate amplified action specificity. Each sample was evaluated in duplicate. Cycle threshold (Ct) of each reaction was automatically calculated within the Rotor-Gene Q system. Delta Ct (ΔCt) was calculated by subtracting the Ct of the housekeeping gene tested (GAPDH) from the gene of interest. ΔCt values of the genes listed above were used to generate a heatmap (Heatmapper ^57^). ΔΔCt of each gene was calculated as the difference between the ΔCt of the differentiated samples and the Day 0 iPSC samples and from these values, relative fold gene expression (2^-(ΔΔCt) was calculated.

#### Immunocytochemistry

At several timepoints along the differentiation spectrum, cells cultured on cover slips were paraformaldehyde-fixed (4%, pH 7.4; PFA) overnight at 4°C. The cover slips were blocked for 2 hours at room temperature with 5% normal donkey serum, 3% bovine serum albumin, 0.1% Triton-X in PBS and then incubated with the primary antibody diluted in this blocking buffer for 6 hours at room temperature. Coverslips underwent 3 washes with blocking buffer followed by incubation with the secondary antibody diluted in blocking buffer for 2 hours at room temperature in the dark. Coverslips were washed with PBS 3 times and sealed to microscope slides (ProLong Diamond Antifade Mountant with and without DAPI) and left to cure for 24 hours at room temperature in the dark. Once properly cured, slides were sealed with nail polish and imaged. The primary antibodies used were: anti-Sox1 (1:150, R&D AF3369), anti-Ki67 (1:100 MAB4190), anti-Sox2 (1:100 Ab5603), anti-TBR2 (1:200 MAB6166), anti-Pax 6 (1:100 901301; BioLegend, San Diego, CA), anti-Tuj1 (801202; BioLegend), anti-Ctip2 (Ab 18465), anti-Nestin (1:400 BD611658), anti-DCX (1:500 SC8066), anti-n-CAM ( 1:100 SC106 lot#C0916), anti-TBR1 (1:100 Ab31940), anti-Stathmin (1:500; ProteinTech Group, Inc., Rosemont, IL), anti-SATB2 (1:400 Ab51502), anti-NeuN (1:100 Millipore ABN78), anti-Stem 123 (1:200 Y40420; Cellartis, Takara Bio Europe AB, Gothenburg, Sweden), anti-NF-H (1:200 Aves NFH7857983), anti-Olig 2 (1:100 Ab9610). CY3, CY5 and Alexa fluorescent conjugated antibodies (Jackson Laboratories) were used as secondary antibodies at dilution of 1:250.

### Animals

#### Strain

We used 6-week-old female athymic RNU rats (n=100; Crl:NIH-Foxn1^rnu^; Charles River Laboratories, Wilmington, MA). These rats were selected for their immunodeficiency, which allows for xenografting without fear of rejection. Based on our personal observations we found that this strain is diurnal, less prone to aggression in social housing, easily trainable, and mild-tempered, thereby making them ideal candidates for intensive behavioral assessments. Rats were housed two per cage in our animal facility under a 12/12 hour light/dark cycle and were provided with food and water *ad libitum*. All animal procedures were performed at Stanford University, under an approved Administrative Panels on Laboratory Animal Care Protocol.

#### Randomization

Each cage (housing two rats) was randomly assigned to an experimental condition group so that both cagemates underwent the same procedures and treatments. Behavioral data was collected and processed by timepoint, and not by experimental group. Tissue data was collected and processed randomly, instead of grouped.

#### Blinding

During transplantation, the surgeon was blinded to the contents of the transplanted material. Rats were given number identifications and trained scientists blinded to the experimental conditions of each individual rat performed all tests and data analyses.

#### Animal mortality and adverse events

Four animals died prior to the final timepoint, and therefore data from these rats were excluded. The most common adverse events were weight loss, epidermal lesions or infections, and ocular irritations - the majority being strain-associated, or social housing-induced nuances. If a rat exhibited significant adverse events, lost more than 5% of its body weight for a sustained amount of time, showed motor deficits not common to the injury model, had no evidence of iPSC-DCNs despite transplantation, or the tissue was unusable, it was excluded from the study (n=26). The mean weight prior to injury was 217.98g ± 4.35, which is within 3% of the mean weight prior to euthanasia, 223.41g ± 7.02.

### Injury Protocol

#### Cervical Hemisection Injury

At eight weeks old, rats (n=76) received a right unilateral hemisection between the 5^th^ and 6^th^ cervical levels (C5/6) as previously described ^58, 59^. Briefly, animals were weighed and placed under inhalation anesthesia using isoflurane (2.5%, 1.5% O_2_) before surgery. A dorsal, midline skin incision was made, and the skin and underlying muscle layers teased apart from the third cervical (C3) segment to the second thoracic (T2) segment (location determined by counting vertebrae and using the distinctive T2 spinous dorsal process as a landmark). The spinous processes were exposed and a C5 dorsal laminectomy performed to expose the underlying spinal cord. Once exposed, the cord was hemisected using a #11 microscalpel (Feather Safety Razor Co., Ltd., Japan) on the right side under microscopic observation, taking care to avoid damaging the anterior spinal artery.

#### Post-operative Care

After injury, the individual muscle layers were sutured (COATED VICRYL^®^ 5-0 (polyglactin 910) Suture; Ethicon, Somerville, NJ), and the skin closed using 9mm wound closure clips. The rat was placed in a temperature-controlled recovery cage until walking and righting abilities were regained. During surgery and twice daily for three days post-operatively, the rats received analgesic (buprenorphine HCL, 0.14 mg/kg), antibiotics (penicillin, 115 mU/kg), and saline. All animals were inspected daily for healing, body condition, and pain with veterinary care given as deemed appropriate.

#### Transplantation Protocol

At two weeks post injury (simulating a sub-acute injury), animals were anaesthetized, reopened, and the injury site exposed as described above. The scar tissue was gently removed under microscopic observation and a 4 μL intralesion micro-injection was given. The injection groups were as follows: i) 150,000 iPSC-DCNs dispersed within a fibrinogen/thrombin matrix with a combination of growth factors ^13^, ii) 150,000 iPSC-DCNs dispersed within a fibrinogen/thrombin matrix without growth factors, or iii) containing no cells and controlling for the insertion and surgery (sham). Rats were then sutured and cared for post-operatively as described above. Growth factors were used at the following final concentrations as previously described by Lu and colleagues (2012): BDNF (50 ng/ul), NT-3 (50 ng/uL), GDNF (10 ng/uL), IGF-1 (10 ng/uL), BFGF (10 ng/uL), aFGF (10 ng/uL), EGF (10 ng/uL), PDGF-AA (10 ng/uL), HGF (10 ng/uL), MDL28170 (50 uM), and VEGF (10 ng/uL) ^13^.

### Animal Behavioral Assessments

#### General

Following transplantation until euthanasia, rats underwent behavioral assessments to monitor changes in forelimb. Cohorts of rats were separated out for additional experimentation prior to sacrifice including anterograde and retrograde neuronal tracing and diphtheria ablation of the transplant. Rats were euthanized from 12 weeks to 52 weeks post transplantation; tissue was excised and analyzed for changes in injury profile, integration, and synaptogenesis.

#### Pre-training and Conditioning

All animals were handled and pre-trained in the following tests for a minimum of two weeks prior to the start of the study. Animals were allowed to acclimate to the behavioral testing facility for two hours prior to testing. Testing occurred at the same time of day and was performed by the same personnel to maintain continuity. The resulting data was analyzed in duplicate by trained researchers blinded to the experimental conditions.

#### Grip Strength Assessment

Grip strength was measured using the TSE Grip Strength Meter as previously described ^60^. Briefly, the rat was allowed to grab the force-metered bar with both forepaws and the subsequent grip force was recorded. Each animal was tested five times per time point with the lowest and highest values discarded and the remaining three values averaged. Initial readings are given in the unit Ponds and were converted to Newtons. Individual grip strengths were averaged per group and represented as a percentage change from pre-injury grip strength force over time. This test was repeated across three sets of animals.

#### Cylinder Forelimb Preference

Forelimb preference was performed as previously described ^58^ with some adjustments as described below. The cylinder test encourages use of the forelimbs for vertical exploration and can be a measure of rat forelimb preference before and after injury. By quantifying the location of forelimb touches as above and below the midline of the rat’s body, the effect of injury and transplantation on balance in conjunction with forelimb use can be measured. Briefly, forelimb preference was measured by placing the rat in a clear Plexiglass cylinder and filming spontaneous exploratory behavior for three minutes. Mirrors were placed at an angle behind the cylinder so that the forelimbs could be viewed at all times and positions. The number of times the rat touched its left or right forepaw to the cylinder above and below the midline of its body was tallied and analyzed for changes in symmetry from pre-injury.

#### Gait Analysis

Gait and locomotion were assessed by tracking footprints as a rat voluntarily traverses a glass beam towards a goal box using the CatWalk^TM^ XT Gait Analysis System (Noldus Information Technology Inc., Leesburg, VA) ^61^. In a darkened room, a camera from beneath the glass beam captured the pressure intensity of the footsteps to file; the data was then uploaded to a program that analyzed the pattern and intensity of the footsteps for contact density and time, swing duration and speed, and stride length. The results were averaged per experimental condition for changes in gait and locomotion.

#### Ladder Rung Test

The ladder rung test measures changes in skilled walking and sensorimotor function ^24^. The testing apparatus consists of a horizontal ladder with removable metal rungs suspended within a clear Plexiglass structure and attached to a “goal” area (where food treats are located) on one end. At each time point, the pattern of ladder rungs was altered to prevent the rat from learning the pattern and anticipating the position of the rungs, thereby standardizing the difficulty of the test. Briefly, rats were filmed walking across the ladder three times per time point. The subsequent videos were watched at half speed to quantify the amount of total right forepaw steps, the number of times the right forepaw completely missed the ladder rung, and the number of times the right forepaw was placed incorrectly (eg. used wrist or digits to balance on the rung, or missed the rung but was able to correct the motion). The percentages of total and partial missteps were calculated per rat and averaged per experimental condition to determine changes in sensorimotor function from baseline over time.

### *In Vivo* Neuronal Tracing

#### Retrograde Neuronal Tracing

Using Fluorogold (Fluorochrome, LLC, Denver, CO) retrograde neuronal tracing was used to determine which ipsi- and contralateral brain cell populations were able to innervate one segment beyond the lesion site. At 12 weeks post transplantation, a subset of rats (n=10) were retrograde traced with Fluorogold (Fluorochrome, 2% solution in sterile distilled H_2_O). Briefly, rats were anesthetized and C5 was exposed as described above. The animal was secured in a stereotactic assemblage and a combined total of 0.6 µL of Fluorogold was pressure injected into three different injection sites, 1.2 mm deep, caudal to the lesion (between the C6/7 vertebrae) at a rate of 100 nL/min followed by a 1 minute rest prior to needle removal. Rats were then sutured and cared for post-operatively as described above. After a 10 day incubation, the rats were euthanized.

#### Anterograde Neuronal Tracing

Anterograde neuronal tracing with Channel Rhodopsin 2 (ChR2; produced by the Stanford University Neuroscience Gene Vector and Virus Core) was used to visualize the projections of the corticospinal tract through the cord in relation to injury and transplantation. At 12 weeks post transplantation, a subset of rats (n=14) were anterograde traced using human channel-rhodopsin (hChR2) coupled to adeno associated viruses (AAV) with either an eYFP reporter protein operating under the CMV promotor (AAV DJ-CaMKIIa hChR2 (E123A)-eYFP-WPRE) or mCherry reporter protein and operating under the hSyn1 promoter (AAV DJ-hSyn1 hChR2(H134R)-mCherry) (Stanford University Neuroscience Gene Vector and Virus Core). Briefly, rats were anesthetized as described above and the skull was stereotactically secured. A dorsal midline incision using a #11 scalpel was made to expose the sutures of the skull and the skin retracted using alligator clamps. To visualize bregma and reduce bleeding, a 1% hydrogen peroxide solution was applied with a cotton swab to the surface of the skull. Four burr holes were drilled over each motor cortex (coordinates using bregma as point 0: 1.5, 1.5; 2.5, 1.0; 2.5, 2.0; 3.5, 1.0) and anesthesia was reduced to 1%. A combined total of 0.8 µL of virus was injected into each cortex at a rate of 75 nL/min at a depth of 1.5 mm with 1 minute rest prior to needle removal. Rats were then sutured and cared for post-operatively as described above. After a 4 week incubation (empirically determined), the rats were euthanized.

### Transplant Ablation

Unnicked diphtheria toxin (DPT; #150; List Biological Laboratories, Inc., Campbell, CA) was used to non-invasively and selectively ablate transplanted human iPSC-DCNs as DPT has a 100,000 fold affinity for human over rat cells ^62^. At 31 weeks post transplantation, a subset of rats (n=10) received two intraperitoneal injections of 50 ug/kg body weight DPT, 24 hours apart ^62^. Rats were euthanized 3 or 7 days after the second injection, with a final behavioral assessment performed prior to euthanasia.

### Perfusion and Tissue Preparation

Animals received an intraperitoneal injection of a pentobarbital and phenytoin solution (VEDCO; 500 mg/kg body weight). Once surgical-plane anesthesia was reached, animals were transcardially perfused with 200 mls of 0.1M PBS followed by 300 mls of paraformaldehyde (4%, pH 7.4; PFA). Following perfusion, brain and spinal cords were dissected and postfixed in PFA for 24 hours at 4°C, and then cryoprotected in a 30% sucrose PBS solution. Tissue segments were embedded in 10% porcine gelatin, postfixed in PFA, cryoprotected in a 30% sucrose PBS solution, and sectioned in coronal, transverse, or sagittal orientations on a freezing microtome with a section thickness of 60 µm.

### Immunohistochemistry

#### Primary Antibodies

The following primary antibodies were used in this study: 5-HT (20080, 1:1000 dilution; Immunostar, Hudson, WI), Ctip2 (ab18465, 1:50 dilution; Abcam, Cambridge, United Kingdom), GFAP (Z033401-2, 1:500 dilution; Dako, Agilent, Santa Clara, CA), GFP (GFP-1020, 1:5000 dilution; Aves Labs Inc., Davis, CA), Homer (160003, 1:200 dilution; Synaptic Systems GmbH, Goettingen, Germany), Iba1 (01919741, 1:500 dilution; FUJIFILM Wako Pure Chemical Corporation, Osaka, Japan), mCherry (600-401-P16, 1:1000 dilution; Rockland Immunochemicals, Inc., Gilbertsville, PA), NCAM (106, 1:500 dilution; Santa Cruz Biotechnology, Inc., Dallas, TX), Neurofilament-H (NFH, 1:1000 dilution; Aves Labs Inc.), Stathmin (11157-1-AP, 1:1000 dilution; ProteinTech Group, Inc., Rosemont, IL), Stem101 (AB-101-U-050, 1:200 dilution; Takara Bio, Inc., Shiga, Japan), Stem121 (AB-121-U-050, 1:1000 dilution; Takara Bio Inc.), Stem123 (AB-123-U-050, 1:1000 dilution; Takara Bio Inc.), TBR1 (ab183032, 1:50 dilution; Abcam), TuJ1 (801202, 1:1000 dilution; BioLegend, San Diego, CA), VGlut1 (135 304, 1:150 dilution; Synaptic Systems) and VGlut2 (135 404, 1:150 dilution; Synaptic Systems).

#### Secondary Antibodies

The following secondary antibodies were used in this study at a 1:800 dilution (Jackson ImmunoResearch Laboratories, Inc., West Grove, PA): Donkey anti chicken AlexaFluor 488 (703-545- 155), Donkey anti mouse AlexaFluor 488 (715-545-151), Donkey anti mouse AlexaFluor 594 (715-585- 151), Donkey anti mouse Cy3 (715-165-150), Donkey anti mouse Cy5 (715-175-150), Donkey anti rabbit AlexaFluor 594 (711-585-152), Donkey anti rabbit Cy3 (711-165-152), and Donkey anti rabbit Cy5 (711-175-152).

Tissue sections were blocked for 2 hours at room temperature with 10% normal donkey serum, 0.1% Triton-X, and a PBS solution and then incubated with the primary antibody diluted in this blocking buffer while rocking for 48 hours at 4°C. Sections underwent 3 washes with blocking buffer followed by incubation with the secondary antibody diluted in blocking buffer for 4 hours at room temperature in the dark. Sections were washed with a phosphate buffer solution 3 times and mounted on microscope slides (ProLong Diamond Antifade Mountant (ThermoFisher Scientific) with and without DAPI) and left to cure for 24 hours at room temperature in the dark. Once properly cured, slides were sealed with nail polish and imaged.

### Imaging and Analysis

#### Equipment and Software

All imaging was performed on a Nikon^®^ C2 Confocal microscope (Nikon^®^, Tokyo, Japan) with a 10x, 20x, 40x, and 60x objective, operated by Nikon NIS-Elements AR acquisition software. Laser intensities and acquisition settings were established for individual channels using optimal LUT settings and applied to entire experimental replicates. NIH ImageJ software ^63^ was used for all analysis, as described below.

#### Delineating Transplants from Host Tissue In Vivo

Human iPSC-DCN transplant survival was determined by serial immuno-labeling spinal cord tissue with green fluorescent protein (GFP; for human transplanted cells tagged with GFP), SC101 (human nuclear marker), or SC121 (human cytoplasmic marker). In our hands, these markers robustly label human iPSC-DCNs with no cross reactivity in host rat cells.

#### Reconstructing and Measuring Lesion Cavity Volume

Lesion cavity volume was defined as the absence of tissue as a result of the injury, and not improper post-mortem tissue handling (determined via GFAP^+^ labeling). The area of remaining damaged tissue was not incorporated into these counts. Transplant volume was defined as the main volume that immunohistochemically tagged transplanted cell bodies (identified by SC101^+^ staining) encompassed at 12 weeks post transplantation. Using ImageJ software, cavity or transplant volume was measured in a series of calibrated, reconstructed images per rat and summed. This number was multiplied by 6 (to account for the 1 in 6 series), 60 µm (to account for tissue section thickness), and 10^-9^ to yield the lesion cavity volume in mm^3^. Volumes were averaged per experimental condition. Data from individual rats were excluded if: (i) the tissue was too damaged to reliably determine cavity volume due to injury versus post-mortem handling, (ii) mortality was less than 12 weeks post transplantation, and (iii) if there was no evidence of iPSC-DCNs transplantation (assessed via GFP or human markers SC121 and SC101 immunofluorescence).

#### Measuring Human iPSC-DCN Transplant Volume

SC101 is a marker of human nuclei, and therefore an effective way to differentiate between human transplanted cells and the rat host cellular microenvironment. Using ImageJ software, a series of calibrated images per rat spanning from the brainstem to the cauda equina of the spinal cord were divided in 500 µm increments from the transplant epicenter and SC101^+^ cells quantified. This number was summed per 500 µm increment and multiplied by 6 (to extrapolate for the 1:6 series). Cell counts were then plotted against distance from the transplant epicenter to determine transplant density and dispersal.

#### Measuring Human iPSC-DCN Axonal Extension

SC121 is an established human cytoplasmic marker that labels processes of human cells. Using ImageJ software, a series of calibrated images per rat spanning from the brainstem to the cauda equina of the spinal cord were divided in 500µm increments from the transplant epicenter. The SC121^+^ axonal processes that were visible at the distance increment lines were quantified and summed per increment. This number was then multiplied by 6 (to extrapolate for the 1:6 series) and plotted against distance from the transplant epicenter to determine axonal growth rostral and caudal to the epicenter.

#### Fluorogold^+^ Cell Counts

A series of brain sections in the coronal plane were imaged using an epifluorescence camera (Nikon). Using ImageJ software, thresholds for pixel intensity and cellular size were established and labeled cells fitting these criteria were automatically quantified in calibrated, oriented images. The Paxinos and Watson Rat Brain Atlas, 5^th^ Edition ^64^ was used to ascertain the correct region of the brain for which the labeled cells were seen. Only fully labelled neurons were counted. Counts within each region were multiplied by 3 (to extrapolate for the 1:3 series) and plotted based on ipsilateral and contralateral sides to determine which propriospinal tracts were innervating the injury and transplant area.

#### Anterograde Tracing Axonal Counts

Using ImageJ software, a series of calibrated images per rat spanning from the brainstem to the cauda equina of the spinal cord were divided in 500 µm increments. The virally infected GFP^+^ and mCherry^+^ axonal processes visible at the distance increment lines were quantified and per increment. This number was then multiplied by 6 (to extrapolate for the 1:6 series) and plotted against distance to determine host corticospinal tract regeneration.

#### Fluorescence Intensity

With some stains in which reliable quantitation of individual cells or processes was not possible, the fluorescence intensity was measured using ImageJ software. In calibrated images in series per rat, ROIs were established and mean gray value (sum of pixel gray values / total number of pixels) was measured.

#### Statistical Analysis

All power analyses and statistics were performed within the laboratory using Sigma Plot v14.5, Microsoft Excel (Microsoft), and the National Institute of Standards and Technology Handbook of Engineering Statistical Methods ^65^. All results were also independently confirmed by the Stanford University Biostatistics Core. All data are presented as the mean ± SEM unless otherwise stated. All data were post-hoc analyzed using a one-way analysis of variance (ANOVA) followed by either Tukey’s procedure, Holm-Bonferonni method, Scheffe’s method, or Welch’s t-test as specified in figure legends. Respective sample size and units are indicated in figure legends. In all statistical analyses, a significance criterion of *P ≤ 0.05, **P < 0.01, ***P < 0.001, and ****P < 0.0001 was used.

## Data Availability

The study data and materials are available from the Corresponding Author on reasonable request.

## Supporting information

Supplementary Data Doulames et al 2021

## Acknowledgments

We would like to acknowledge and thank Dr. Theo Palmer (scientific advising, writing assistance, technical editing), Christine Plant (writing assistance, technical editing, language editing, and proofreading), Nnaoma Oji (data collection), Gabe Weininger (data collection), Madhuri Vangipuram (data collection), and Meghan Hefferon (data collection). The work described herein was supported by the: Department of Defense (DoD) W81XWH-18-1-026, National Institutes of Health (NIH) 1R01EB02766602, Wings for Life Spinal Cord Research Foundation WFL-US-15/17, California Institute of Regenerative Medicine (CIRM) RT3-07948, International Spinal Research Trust (United Kingdom) STR119, Wings for Life Spinal Cord Research Foundation (Austria) WFL-US-020/14, Stanford University Bio-X Program Interdisciplinary Initiatives Seed Grant IIP-7, Dennis Chan Foundation, Klein Family Fund, Lucile Packard Foundation for Children’s Health, Stanford Institute for Neuro-Innovation and Translational Neurosciences (SINTN), Saunders Family Neuroscience Fund, James Doty Neurosurgery Fund, Hearst Neuroscience Fund, and Eileen Bond Research Fund.

## Author Contributions

GWP, VMD, and JW were all responsible for conceptualization, investigation, project administration and supervision, methodology, data curation, and formal analysis. VMD and GWP were responsible for visualization, writing, reviewing, and editing. Additionally, GWP was responsible for funding acquisition and resources.

