## Supplementary Data Doulames et al 2021 for "Human Deep Cortical Neurons Promote Regeneration and Recovery After Cervical Spinal Cord Injury"

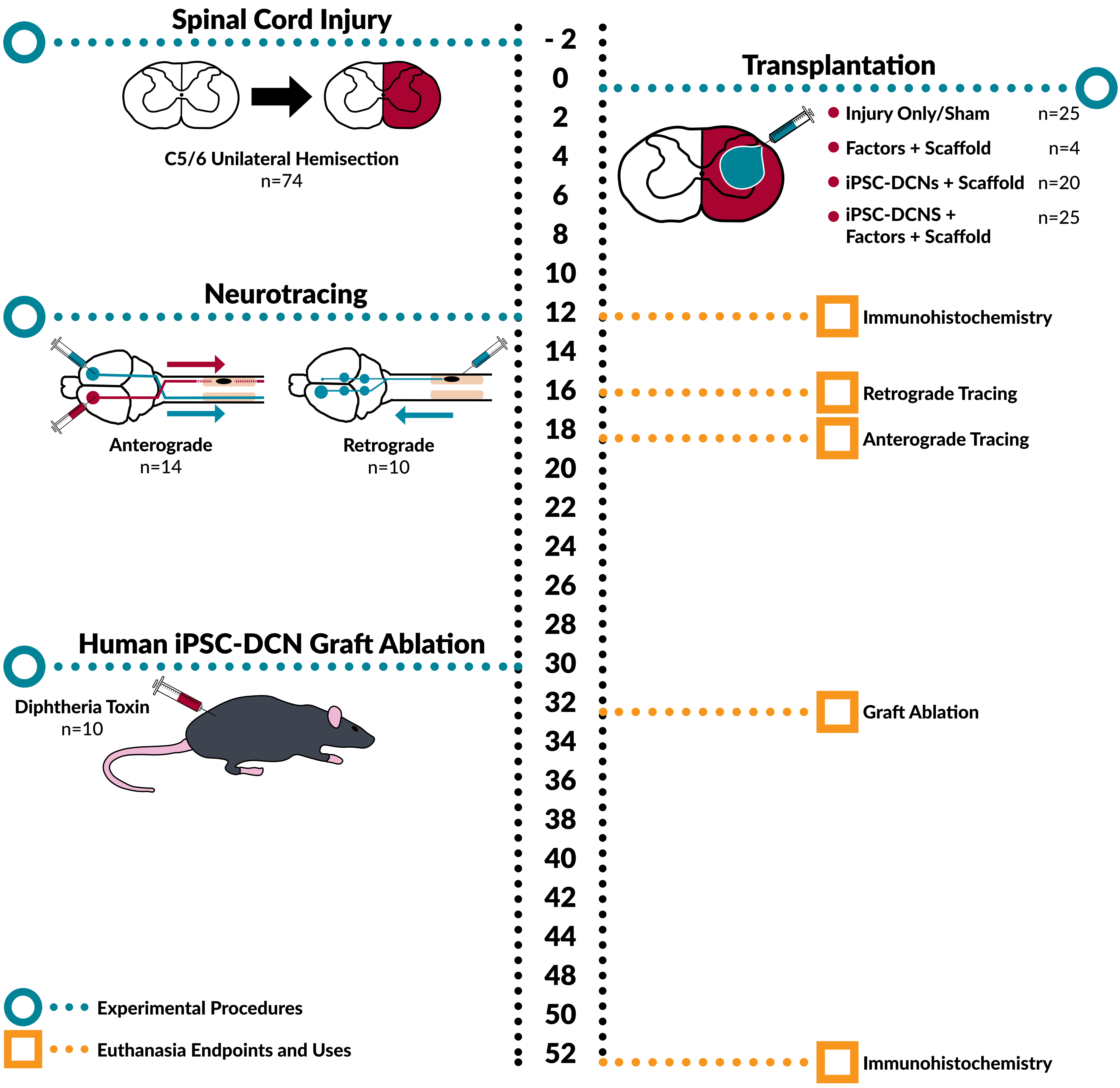

Spinal Cord Injury
C5/6 Unilateral Hemisection
n=74
Neurotracing
Anterograde
n=14
Retrograde
n=10
Human iPSC-DCN Graft Ablation
Diphtheria Toxin
n=10
Experimental Procedures
Euthanasia Endpoints and Uses
- 2
0
2
4
6
8
10
12
14
16
18
20
22
24
26
28
30
32
34
36
38
40
42
44
48
50
52
Transplantation
Injury Only/Sham n=25
Factors + Scaffold n=4
iPSC-DCNs + Scaffold n=20
iPSC-DCNS + Factors + Scaffold n=25
Immunohistochemistry
Retrograde Tracing
Anterograde Tracing
Graft Ablation
Immunohistochemistry

**Fig. S1.**

*Experimental timeline* in vivo. Rats are injured and undergo transplantation 2 weeks later. Subsets of rats undergo anterograde or retrograde tracing at 12 weeks post transplantation. A subset of rats receives Diphtheria Toxin at 30 weeks post transplantation to selectively ablate the human iPSC-DCN graft. Animals are euthanized at 12, 16, 18, 32, and 52 weeks post transplantation.

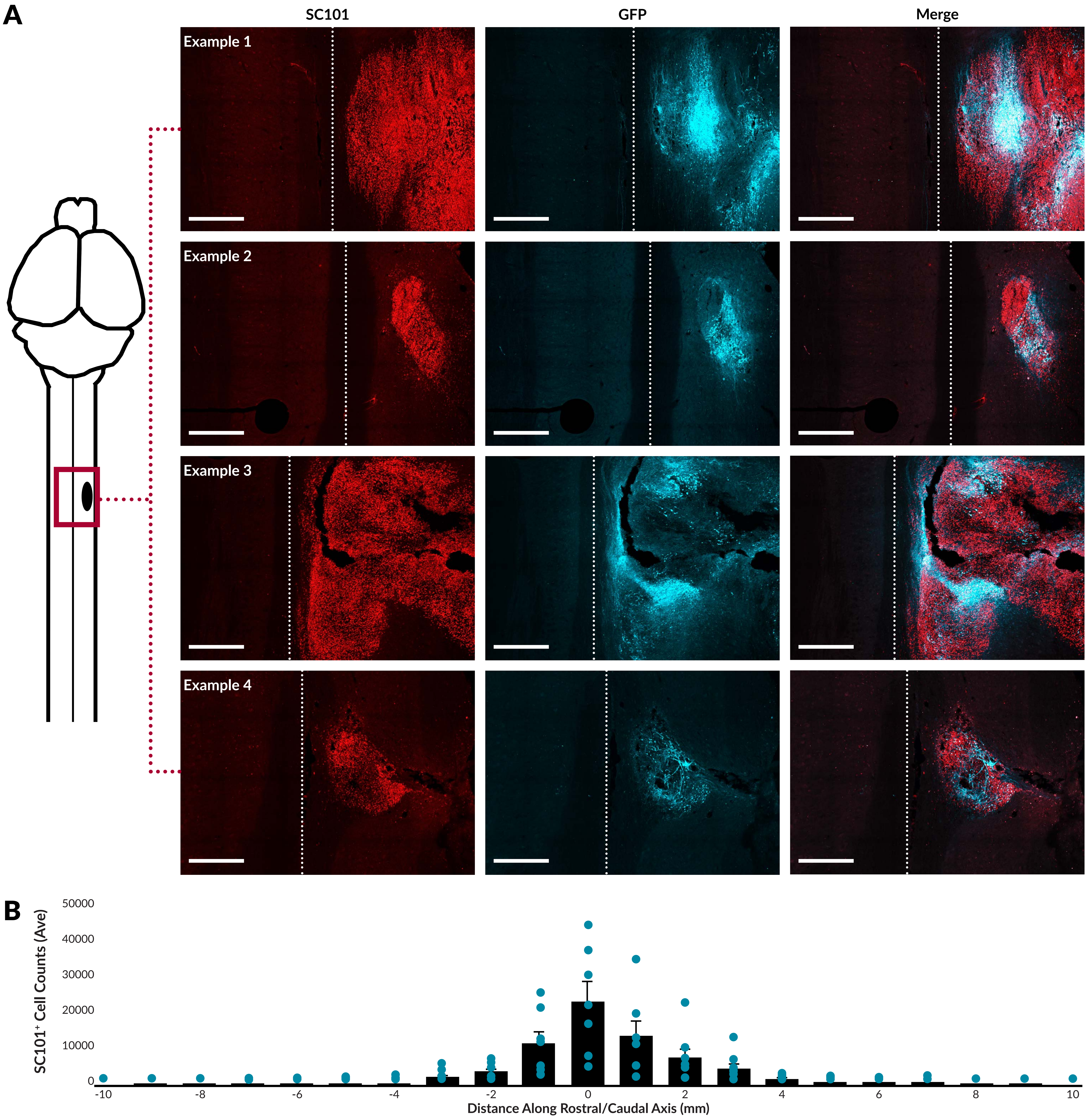
A
SC101
GFP
Merge
Example 1
Example 2
Example 3
Example 4
B
50000
40000
30000
20000
10000
0
SC101⁺ Cell Counts (Ave)
-10
-8
-6
-4
-2
0
2
4
6
8
10
Distance Along Rostral/Caudal Axis (mm)

**Fig. S2.**

*Variability in graft size and cell number*. **A.** Qualitative images depicting variability in SC101$^+$/GFP$^+$ iPSC-DCN graft sizes and numbers at 12 post transplantation. Dotted line indicates center of the spinal cord in transverse plane, as referenced in schematic. Images were taken at 20x magnification in 10 um Z-stacks and compressed into a maximum projection. Images were false-colored and manipulated identically to visually delineate graft from surrounding spinal cord tissue. Scale bars = 500 um. **B.** SC101$^+$ grafted human cells were quantified in 1 mm increments from the graft epicenter after 12 weeks to determine migration and dispersal. Averages are represented as columns ± SEM with individual values overlayed (blue circles).

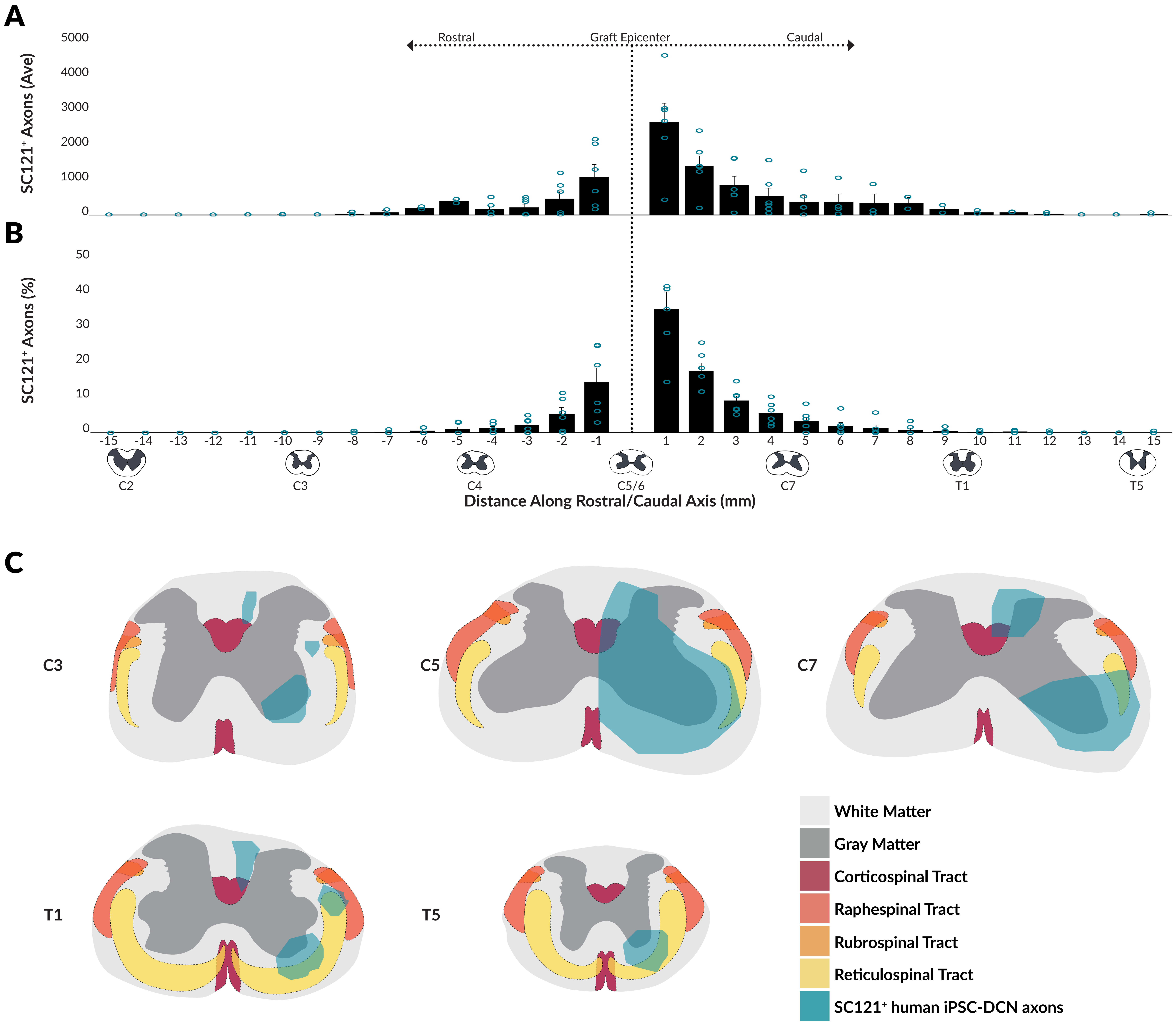
A
Rostral
Graft Epicenter
Caudal
5000
4000
3000
2000
1000
0
SC121⁺ Axons (Ave)
B
50
40
30
20
10
0
SC121⁺ Axons (%)
-15 -14 -13 -12 -11 -10 -9 -8 -7 -6 -5 -4 -3 -2 -1 1 2 3 4 5 6 7 8 9 10 11 12 13 14 15
C2
C3
C4
C5/6
C7
T1
T5
Distance Along Rostral/Caudal Axis (mm)
C
C3
C5
C7
T1
T5
White Matter
Gray Matter
Corticospinal Tract
Raphespinal Tract
Rubrospinal Tract
Reticulospinal Tract
SC121⁺ human iPSC-DCN axons

**Fig. S3.**

*Distribution and span of iPSC-DCN axonal growth*. SC121$^+$ labeled axons of transplanted iPSC-DCNs in a subset of rats (n=8) were quantified in 1 mm increments from the graft epicenter. **A.** Axon number averages are represented as columns ± SEM with individual values overlayed (blue circles). **B.** Axon percentages are represented as columns ± SEM with individual values overlayed (blue circles). **C.** Locations of quantified SC121$^+$ human iPSC-DCN axons throughout cervical to mid thoracic spinal segments are overlayed approximated locations of descending neural tracts.

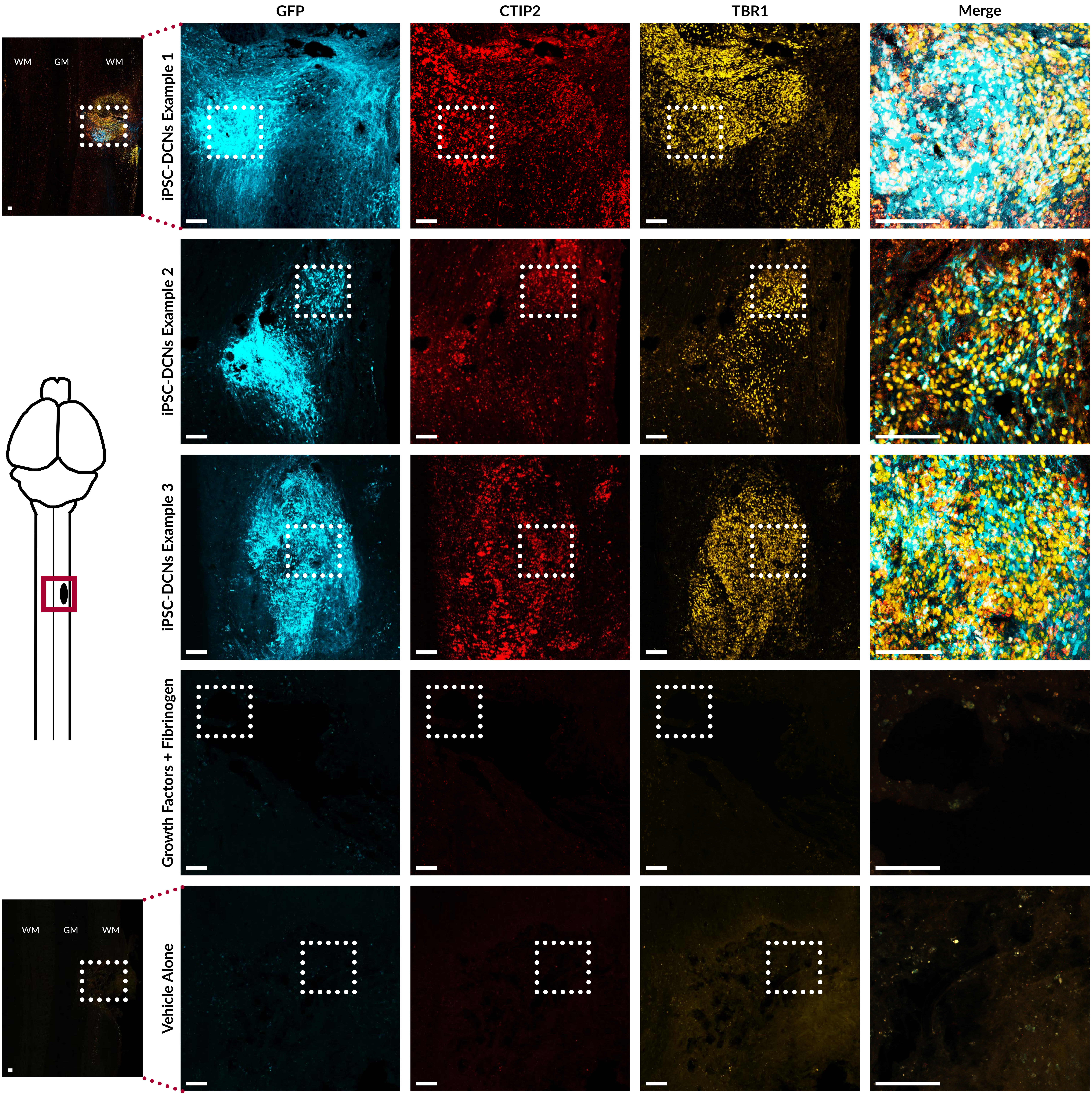
GFP
CTIP2
TBR1
Merge
WM
GM
WM
iPSC-DCNs Example 1
iPSC-DCNs Example 2
iPSC-DCNs Example 3
Growth Factors + Fibrinogen
WM
GM
WM
Vehicle Alone

**Fig. S4.**

*Human iPSC-DCNs express Ctip2 and TBR1*. Qualitative images from 2 different rats depicting colocalization of $GFP^+$ iPSC-DCN grafts with deep cortical markers Ctip2 and TBR1 at 12 post transplantation. Non-cellular controls (growth factors and fibrinogen or vehicle alone) are present to demonstrate the specificity of the GFP, Ctip2, and TBR1 antibodies. Images were taken at 20x magnification in 10 um Z-stacks and compressed into a maximum projection. Images were false-colored and manipulated identically. WM = white matter, GM = gray matter. Scale bars = 250 um.

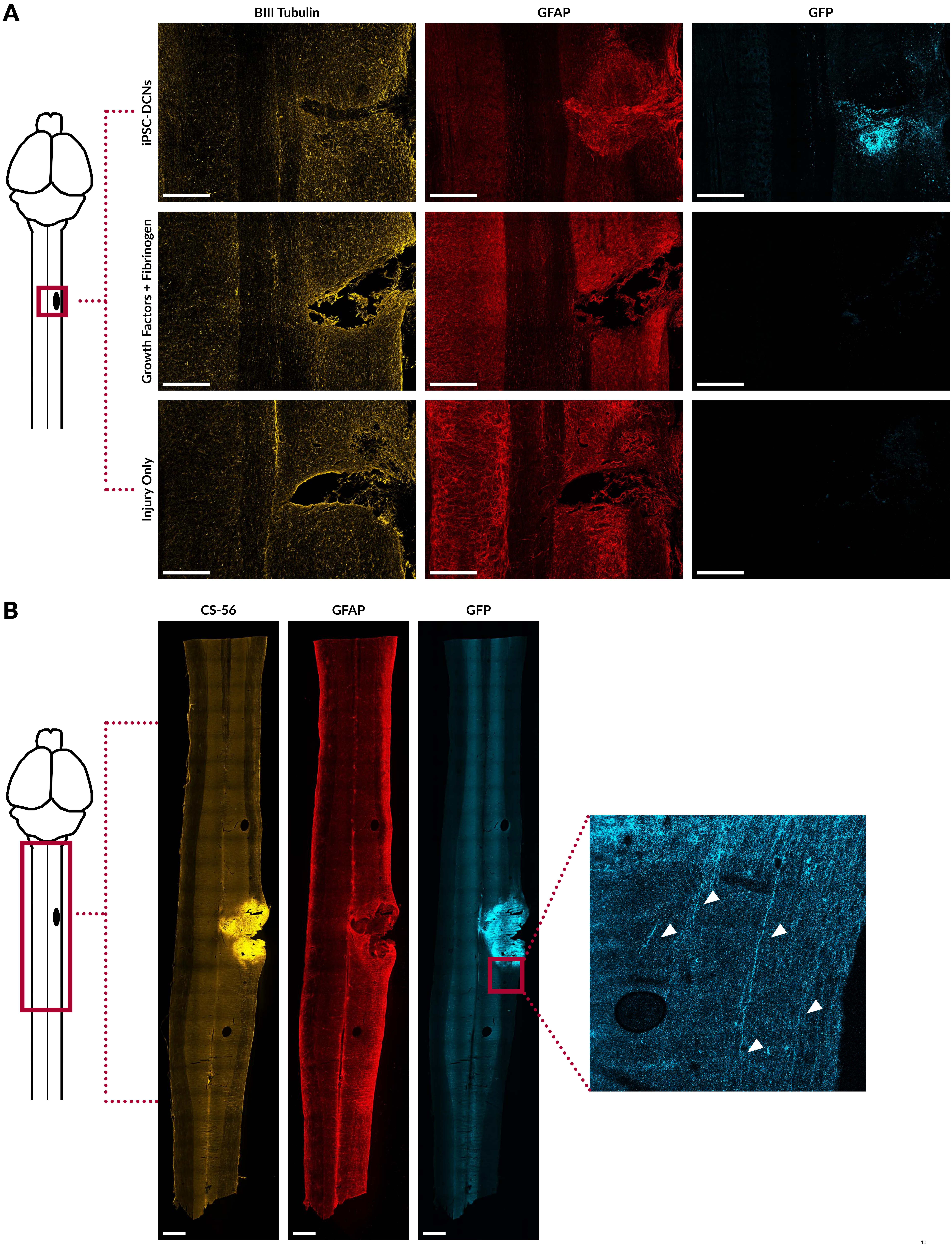
A
BIII Tubulin
GFAP
GFP
iPSC-DCNs
Growth Factors + Fibrinogen
Injury Only
B
CS-56
GFAP
GFP

**Fig. S5.**

*Human iPSC-DCNs ignore components of the SCI scar*. A. Qualitative images depicting bIII tubulin, GFAP, and GFP staining in injured rats transplanted with iPSC-DCNs, growth factors and fibrinogen, or vehicle alone at 12 weeks post transplantation. Images were taken at 20x magnification in 10 um Z-stacks and compressed into a maximum projection. Images were false-colored and manipulated identically. Scale bars = 500 um. B. Qualitative images depicting CS-56, GFAP, and GFP staining in an injured rat transplanted with iPSC-DCNs. Insert shows $GFP^+$ iPSC-DCN axons (white arrows) exiting the lesion/transplantation site. Images were taken at 10x magnification in 10 um Z-stacks and compressed into a maximum projection. Images were false-colored and manipulated identically. Scale bars = 500 um.

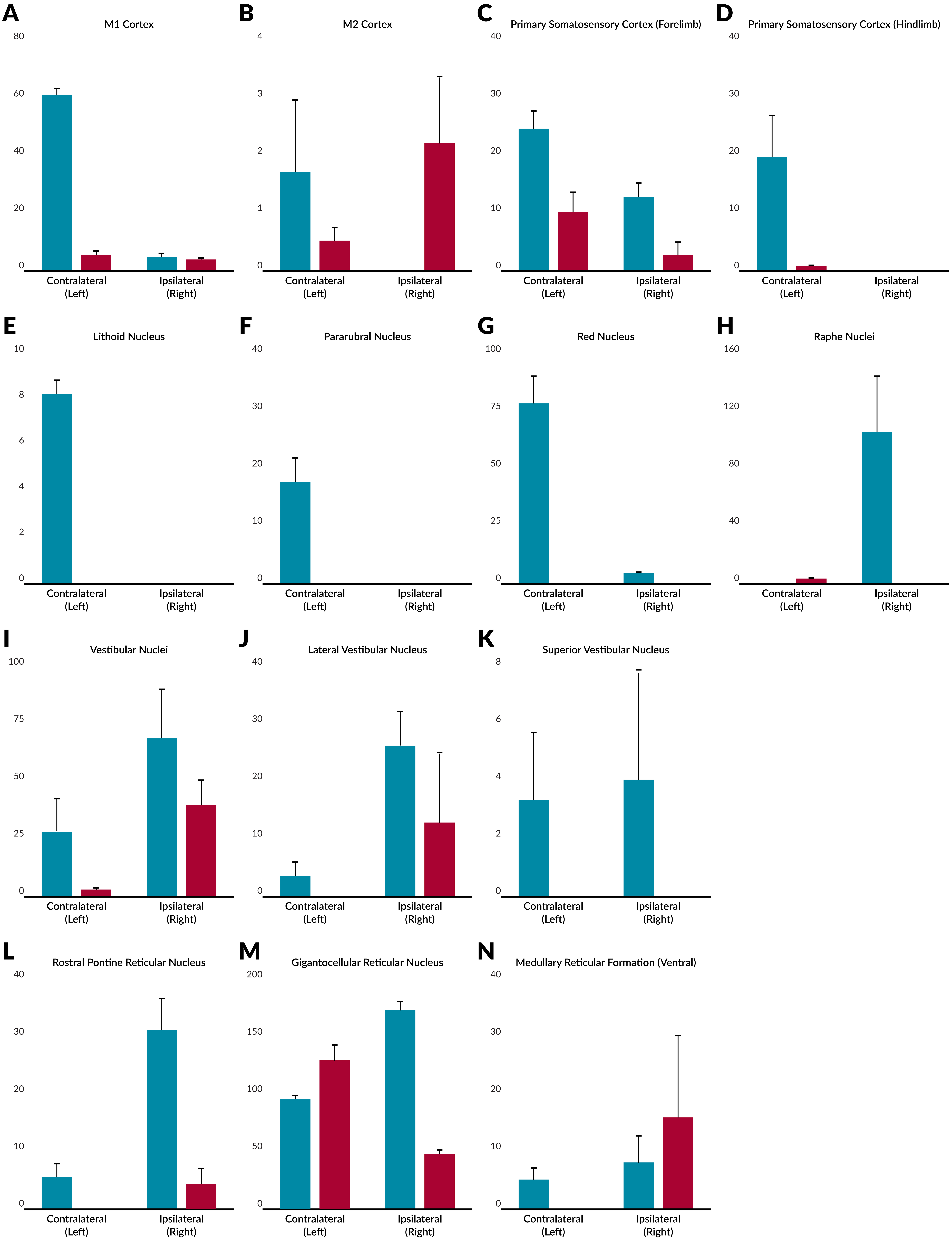
A
M1 Cortex
B
M2 Cortex
C
Primary Somatosensory Cortex (Forelimb)
D
Primary Somatosensory Cortex (Hindlimb)
E
Lithoid Nucleus
F
Pararubral Nucleus
G
Red Nucleus
H
Raphe Nuclei
I
Vestibular Nuclei
J
Lateral Vestibular Nucleus
K
Superior Vestibular Nucleus
L
Rostral Pontine Reticular Nucleus
M
Gigantocellular Reticular Nucleus
N
Medullary Reticular Formation (Ventral)
Contralateral (Left)
Ipsilateral (Right)

**Fig. S6.**

*Human iPSC-DCNs promote supraspinal regeneration*. A.-N. Quantification of Fluorogold$^{+}$ cells within left (contralateral) and right (ipsilateral) brain regions of injured and transplanted rats following retrograde tracing caudal to the SCI. Results are represented as the quantified number of cells averaged across the cohort ± SEM (red = injury control, blue = transplant; n=10 per cohort). Tukey HSD Test; $^{**}P \leq 0.01$, $^{***}P \leq 0.001$.

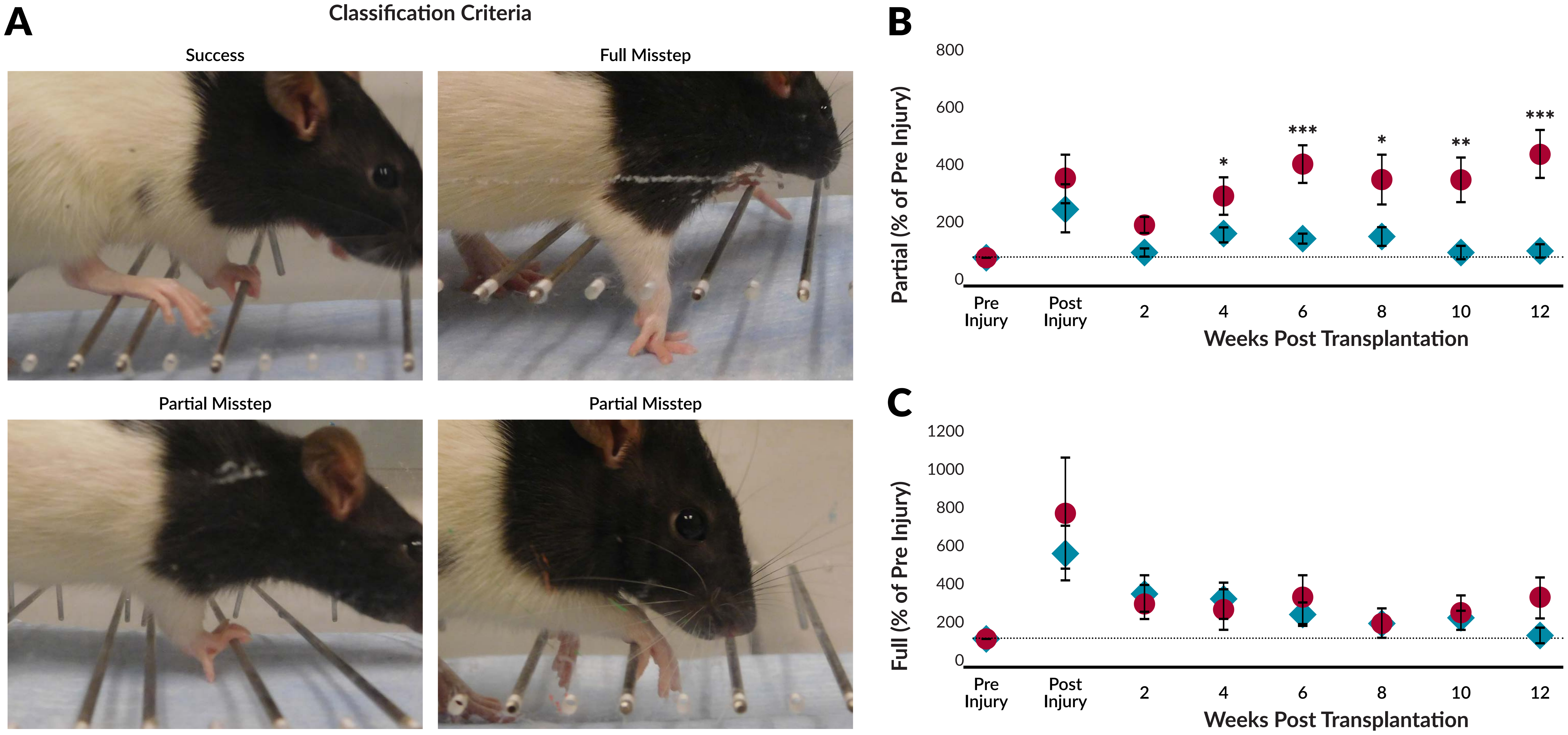
A
Classification Criteria
Success
Full Misstep
Partial Misstep
Partial Misstep
B
Partial (% of Pre Injury)
800
600
400
200
0
***
*
***
*
**
**
Pre Injury
Post Injury
2
4
6
8
10
12
Weeks Post Transplantation
C
Full (% of Pre Injury)
1200
1000
800
600
400
200
0
Weeks Post Transplantation

**Fig. S7.**

*Human iPSC-DCN transplantation differentially affects skilled walking*. **A.** Skilled walking was measured using the ladder rung test. Scoring of the right paw was calculated by quantifying the percentage of instances in which rats successfully grasped the bar, completely missed the bar (full misstep), or grasped the bar using incorrect posture (partial misstep). Representative images were created using age- and size-matched female uninjured RNU+/- rats but all quantification was done in our experimental cohorts (injury control, n=6; and transplanted rats, n=10). **B.** Partial errors as a percentage of pre-injury errors are represented as points ± SEM (red circles = injury control, blue diamonds = transplant). Welch's T-Test for pairwise comparisons between cohorts at each timepoint; $*P \leq 0.05$, $**P \leq 0.01$, $***P \leq 0.001$. Dotted line denotes pre-injury baseline. **C.** Full errors as a percentage of pre-injury errors are represented as points ± SEM (red circles = injury control, blue diamonds = transplant). Welch's T-Test for pairwise comparisons between cohorts at each timepoint; all were nonsignificant. Dotted line denotes pre-injury baseline.

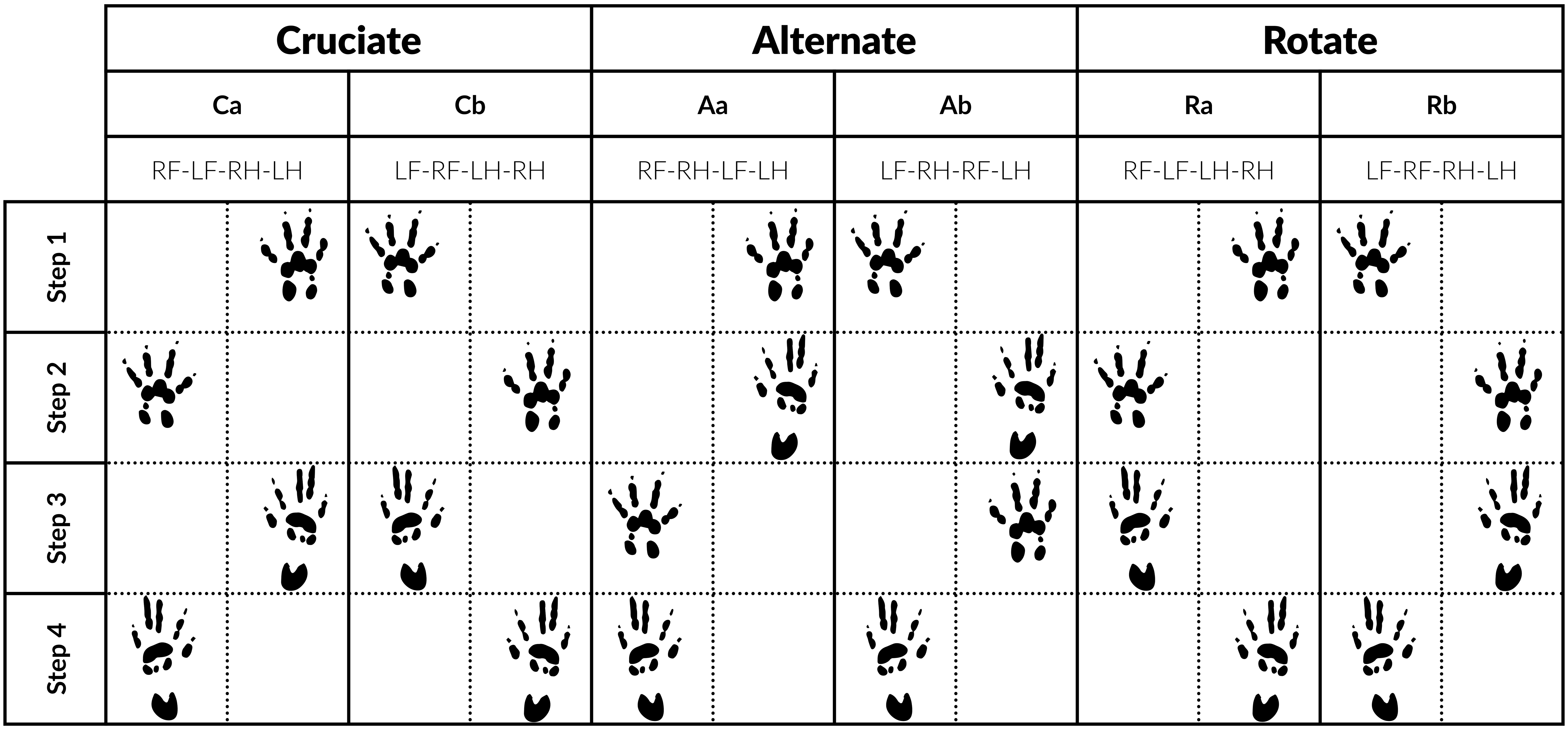
Cruciate
Alternate
Rotate
Ca
Cb
Aa
Ab
Ra
Rb
RF-LF-RH-LH
LF-RF-LH-RH
RF-RH-LF-LH
LF-RH-RF-LH
RF-LF-LH-RH
LF-RF-RH-LH
Step 1
Step 2
Step 3
Step 4

**Fig. S8.**

*Rodent locomotion patterns*. Rodents have 6 major locomotion patterns that fall into 3 groups. Of these, the AB pattern is most prevalent in uninjured animals.
